# A screen for adherens junction proteins regulating collective cell migration and testis morphogenesis reveals important roles for the Rab GAP RN-tre and the kinase Par-1

**DOI:** 10.64898/2026.05.22.727264

**Authors:** Sarah E. Clark, Siobhan C. Morris, Joseph B. Dordor, Lawrencia S. Amo, Renick Wiltshire, Taino Encarnacion, Maik C. Bischoff, Mark Peifer

## Abstract

Animal tissues have diverse architectures and cell behaviors across the epithelial-mesenchymal spectrum. Cell adhesion mediated by classical cadherins is foundational. Cadherins nucleate complexes of dozens of proteins connecting junctions to the cytoskeleton and signaling downstream. Many junctional proteins are well-studied in epithelia, but less is known about roles during mesenchymal migration. The nascent myotubes of the pupal Drosophila testis provide an excellent model for N-cadherin mediated mesenchymal migration. We combined a proximity proteomics dataset of adherens junction proteins in mammalian epithelial cells with genome-wide shRNA libraries knocking down Drosophila genes to begin to define the subset of junctional proteins important in mesenchymal migration. While N-cadherin is predominant, E-cadherin plays a supporting role. Surprisingly, several proteins with key roles in epithelial morphogenesis, including Afadin’s homolog Canoe, ZO-1’s homolog Polychaetoid, and Par3’s homolog Bazooka play at most modest roles. Twenty-two genes with diverse cell biological roles had strong to moderate defects in testis morphogenesis. These will provide a community resource. We followed up two. The kinase Par-1 is important for migration and gap closure, with knockdown phenotypes paralleling those of myosin. The Rab GAP RN-tre does not have roles until after migration and works in parallel with N-cadherin during testis spiralization.

## Introduction

The array of organs in the animal body differ dramatically in their cellular architecture, but cell-cell adhesion mediated by cadherins is a common principle. The cadherin protein family diversified in different animal phyla, allowing family members to carry out functions ranging from mediating tissue architecture to polarizing tissues along the body axes to wiring synaptic connectivity in the nervous system (Oda and Takeichi, 2011). Classical cadherins were the first family members discovered and are present in all animals. All share cadherin repeat domains in their extracellular region that mediate homophylic adhesion, and conserved sequences in the cytoplasmic tails bind conserved catenins. These proteins help define tissue architecture and organ function in animals ranging from cnidarians (Nathaniel Clarke *et al*., 2019) to flies (Tepass *et al*., 1996; Uemura *et al*., 1996) and mammals (Larue *et al*., 1994; Riethmacher *et al*., 1995).

Classical cadherin functions have been studied most intensively in vertebrates and Drosophila, where they mediate diverse tissue architectures and cell behaviors. E-cadherin was the first cadherin identified and it is expressed in most epithelial tissues (Yoshida-Noro *et al*., 1984), where it plays essential roles from embryo compaction onward (Takeichi, 2014). The second classical cadherin identified, N-cadherin, has a different pattern of expression (Hatta and Takeichi, 1986). It is expressed in neural lineages from neural tube invagination to differentiation as neurons, and in multiple mesodermal derivatives, both during collective migration and in at least a subset of differentiated tissues like skeletal muscle and the heart (Radice, 2013). Intriguingly, an independent gene duplication in the insect lineage led to cadherins whose expression patterns largely mirror those of Ecad and Ncad. In Drosophila, DE-cadherin (encoded by the *shotgun* gene; henceforth Ecad) is expressed in essentially all epithelial tissues (Tepass *et al*., 1996; Uemura *et al*., 1996) while DN-cadherin (henceforth Ncad) is expressed in the CNS and in many mesodermal derivatives (Iwai *et al*., 1997), both as they migrate throughout the body and in differentiated tissues like muscle.

In addition to mediating cell-cell adhesion, classical cadherins also connect cell junctions to the actomyosin cytoskeleton, thus regulating cell shape change and cell rearrangement in embryonic and post-embryonic tissues (Campas *et al*., 2024). This linkage was initially thought to be simple, mediated by a linear chain of interactions from the cadherin tail through beta-catenin to α-catenin to actin, but we now know it is much more complex. Dozens of proteins assemble at epithelial cell-cell adherens junctions (AJs) via multivalent linkages, providing both robustness and mechano-responsiveness to the cytoskeletal linkage (Perez-Vale and Peifer, 2020). Functional analysis revealed that individual proteins vary in their importance in epithelial morphogenesis. The core cadherin-catenin complex is essential for adhesion itself. Proteins like Canoe (Cno) and Polychaetoid (Pyd) and their mammalian orthologs Afadin and ZO-1 are dispensable for adhesion but play essential roles in strengthening junctions under mechanical tension, rendering them critical for many morphogenetic movements. In contrast, other proteins like Ajuba (Jub) and Sidekick play supporting roles: null mutants are viable but close inspection reveals defects in cell arrangements and junctional stability.

One key question for the field is how different tissues, all relying on classical cadherins, adopt different cell architectures and behaviors. In Drosophila, tissues expressing Ecad usually have epithelial architecture, even in tissues where cells are rearranging, while Ncad is often predominant in migrating mesenchymal tissues. The classical epithelial-to-mesenchymal transition in mammals is pictured as involving downregulation of Ecad and upregulation of Ncad (Loh *et al*., 2019), but it is now clear that events like metastasis of tumors in epithelial tissues can occur without downregulation of Ecad (Shamir and Ewald, 2015). However, these are also tissues in which both Ecad and Ncad play roles, such as the developing pupal eye (Hayashi and Carthew, 2004). Despite decades of work, it remains unclear which features of cell-cell junctions lead to the diverse behaviors seen along the epithelial-mesenchymal axis. A variety of mechanisms have been suggested (Wheelock *et al*., 2008), which may operate in parallel. These include regulation by E- or N-cad of distinct receptor tyrosine kinases (e.g. (Williams *et al*., 1994; Qian *et al*., 2004), interactions with different isoforms of p120catenin (Seidel *et al*., 2004), and differential effects on Wnt-signaling (Libusova *et al*., 2010). Another potential mechanism involves the use of different complexes of junction:cytoskeletal linker proteins to differentially alter the actomyosin cytoskeleton and thus cell behavior (Wheelock *et al*., 2008; Loh *et al*., 2019).

To test the latter hypothesis, we first needed a list of the proteins that assemble at epithelial AJs. We and others used proximity proteomics to identify proteins that physically localize to mammalian epithelial AJs, using either Afadin (Baskaran *et al*., 2021; Goudreault *et al*., 2022; Choi *et al*., 2025) or Ecad (Guo *et al*., 2014; Van Itallie *et al*., 2014) as the starting point. In our work we fused a promiscuous biotin ligase, BirA, to each end of mammalian Afadin, and expressed the fusions in canine MDCK cells, a premier epithelial cell model (Choi *et al*., 2025). We curated a list of 144 validated proteins, a list that substantially overlapped similar lists generated by others. Many of these proteins have known roles in epithelial cell junctions, though others are less well studied.

We and others have explored the function of many of these proteins in Drosophila epithelial tissues (Perez-Vale and Peifer, 2020), including the embryonic ectoderm, the imaginal discs, and the follicular epithelium. However, most of these proteins have not been functionally characterized during mesenchymal migration, in tissues where Ncad is thought to be the primary player. Here we use our Ecad-based proteomic resource to begin to define whether migrating mesenchymal cells use the same subsets of junctional proteins to regulate cell behavior. As our model for Ncad-mediated mesenchymal cell migration, we use the nascent myotubes of the pupal *Drosophila* testis (Bischoff and Bogdan, 2021). These myotubes migrate from the genital disc onto the testis during pupal development, ∼31 hours after pupariation (APF) (Fig 1A), ultimately fully surrounding the testis and differentiating into a circumferential sheet of muscles. The testis subsequently re-shapes itself into the spiral seen in the adult (Fig 1B). Testis myotubes express Ncad during and after migration, and Ncad knockdown leads to defects during migration and to gaps in muscle adhesion along the proximal-distal axis in the adult (Fig 1A vs D-G; (Rothenbusch-Fender *et al*., 2017; Bischoff and Bogdan, 2021). To begin to define which junctional proteins are essential in this tissue, we identified Drosophila homologs of the top hits from our mammalian epithelial cell proximity proteomics screen and used shRNAs to knock them down in a tissue-specific way in the mesenchymal myotube cells during migration.

**Fig 1.**
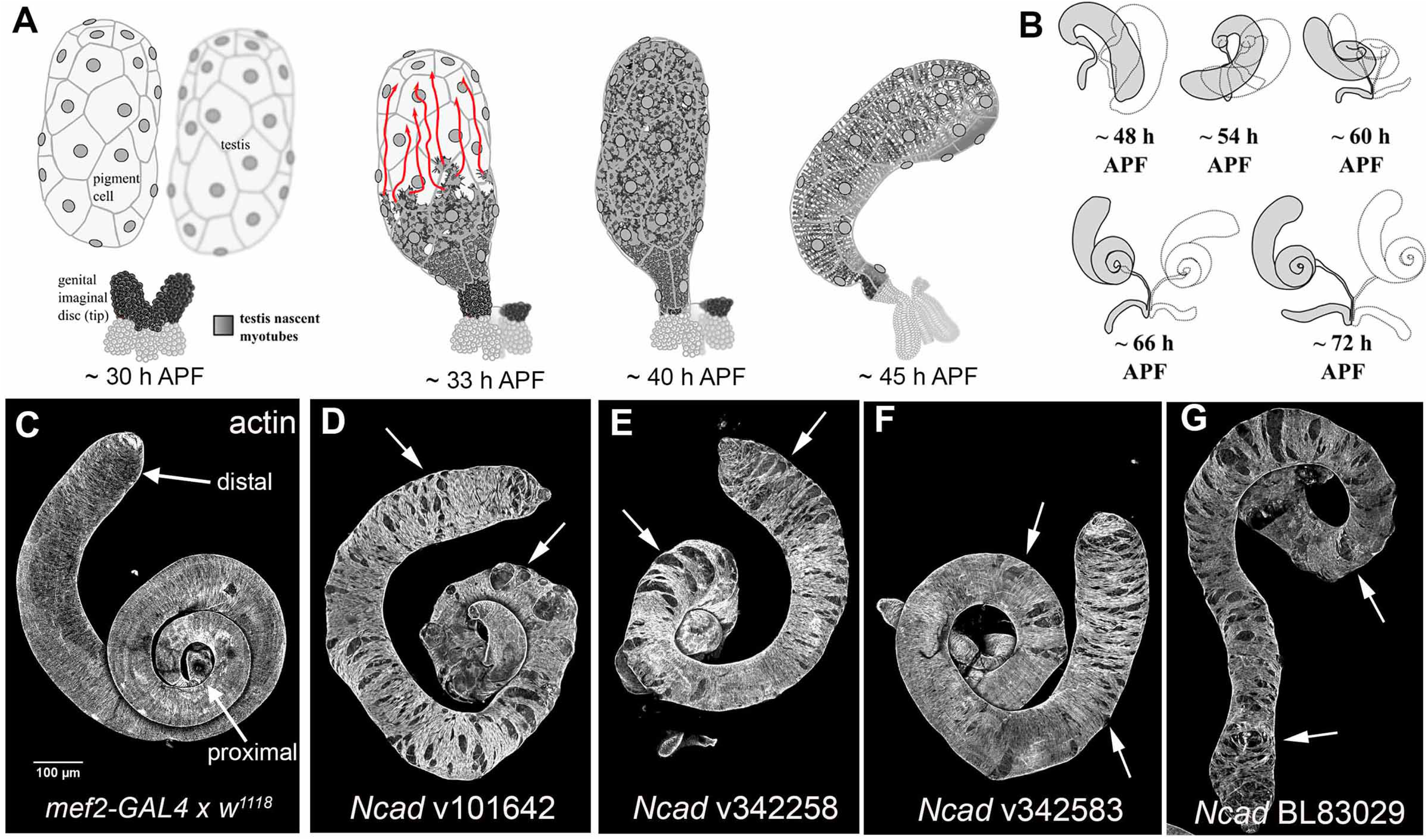
N-cadherin is an essential regulator of testis nascent myotube migration. **(A)** Diagram illustrating testis nascent myotubes collective cell migration during pupal *Drosophila* testis development (derived from (Bischoff and Bogdan, 2021). Myotubes begin on the genital disc, and by 31-32 hours After Puparium Formation (APF) have started to migrate onto the testis, into the narrow space between underlying germline cells and overlying pigment cells. They migrate from the proximal to the distal end, completing migration by 40 hours and then beginning to differentiate as circumferential muscles. Testis spiralization starts ∼48 h APF. **(B)** Timeline of post migratory shaping of the pupal *Drosophila* testis. **(C-G)** Testes from wildtype adults, and adults expressing knockdowns of Ncad. All are stained with fluorescently-labeled phalloidin to reveal F-actin, highlighting muscles. (C) Wildtype (*mef2-Gal4* crossed with *w^1118^* flies). The wildtype testis is covered completely by circumferential muscle and has a spiral shape. Arrows indicate proximal and distal testis. (D-G) Four Ncad shRNA lines crossed to *mef2-Gal4.* Adult testes have gaps all along the proximal-distal axis (arrows). D-G are included in our Class 2 (cell adhesion defects).

## Results

### Identifying candidate junctional proteins

One means by which classical cadherins could shape different tissue architectures and cell behaviors is by using different subsets of the proteins that link them to the cytoskeleton and transmit adhesion signaling. Our proximity proteomics approach to define proteins localized to mammalian epithelial AJs (Choi *et al*., 2025), used Afadin, the Cno homolog, fused to a biotin-ligase at either its N- or C-terminus, yielding two lists. We first combined these lists, selecting the top 124 ordered by average enrichment over controls, Next, we removed duplicates, as many proteins were identified with both baits (Suppl Table 1). We also eliminated 22 proteins whose known functions made it unlikely they play specific roles at AJs (e.g., the citrate transporter SLC25A1 or Eukaryotic translation initiation factor 4B; highlighted in orange).

With this list in hand, we determined whether each had an apparent Drosophila ortholog. Some had no apparent homolog as assessed by blast searching and consulting the “Ortholog” tab in Flybase. In several cases multiple mammalian family members (e.g. ZO-1, ZO-2, and ZO-3; Nectin 2 and 3; or Scribble and Erbin) were independently identified in our proteomics screen, but Drosophila has only a single family member. This left us with 58 candidates (Table 1). To these we added a small number of genes encoding junctional proteins not identified in our proteomics screen. These included the three Drosophila classical cadherins, three known junctional-cytoskeletal linkers or polarity proteins ( (Armadillo/beta-catenin; Arm), Jub/Ajuba, and aPKC, along with Stepke, the protein partner of Sstn which was included on our candidate list.

Our earlier RNAseq on myotube migration (Bischoff *et al*., 2025b) provided us with a list of genes expressed during migration (31 hr after pupariation (APF)) or after migration was completed and muscles were beginning to elongate and differentiate (45 hr APF; Table 1). Only two genes on the list, *lasp* and *cadN2*, the second Ncad relative, were detected at such low levels that we think it is likely they are not expressed in myotubes at this time point. While RPKM (Reads per kilobase of transcript per Million mapped reads) data cannot provide detailed information about relative expression levels, we did note some trends in our six replicates at each timepoint. During migration (31 hours APF), the RPKM values Ecad were consistently lower than those of *Ncad* in all replicates, and the lowest value for *Ncad* was higher than the highest value for *Ecad* (Suppl Fig 1). Further, the RPKM levels in myotubes of three junctional proteins with key roles in epithelia, *cno, pyd,* and *baz* (Drosophila Par3) were more consistent with those of *Ecad,* and once again the highest value of any of these three genes was lower than the lowest value for *Ncad* (Suppl Fig 1).

### The core catenin Arm co-localizes with Ncad in migrating myotubes but Cno and Pyd may not

We previously found that while Ncad localizes to AJs of migrating myotubes, Ecad is not detectable there, though it does localize to junctions of overlaying pigment cells (Rothenbusch-Fender *et al*., 2017; Bischoff *et al*., 2025a). We thus compared the localization in migrating myotubes of Ncad and the core catenin Arm with those of two key epithelial junctional proteins that work together with Ecad, Cno and Pyd. We used *mef2*-GAL4 driven Lifeact-eGFP and a nuclear-localized mCherry to visualize actin and nuclei specifically in myotubes. Ncad localizes to cell junctions between migrating myotubes (Fig 2A, A”, arrows), as does Arm (Fig 2C, C”, arrows). In contrast, there was no apparent localization of Cno at cell junctions in migrating myotubes (Fig 2A, A”’, arrows; Fig 2C, C”’, arrows), while in the same samples both Arm and Cno exhibited strong localization to the epithelial cells of the seminal vesicle, which is adjacent to the testis and serves to store sperm (Fig 2B, B”,B”’ cyan arrows). We carried out similar imaging of Pyd localization. As with Cno, Pyd did not accumulate at myotube cell junctions (Fig 2D, D”, D”’ arrows), while Pyd accumulated at high levels at cell junctions in the adjacent seminal vesicle, which does not express Ncad (Fig 2E, E”, E”’ cyan arrows). These data suggest that migrating myotubes do not express high levels of some AJ-cytoskeletal linkers with important roles in epithelial tissues.

**Fig 2.**
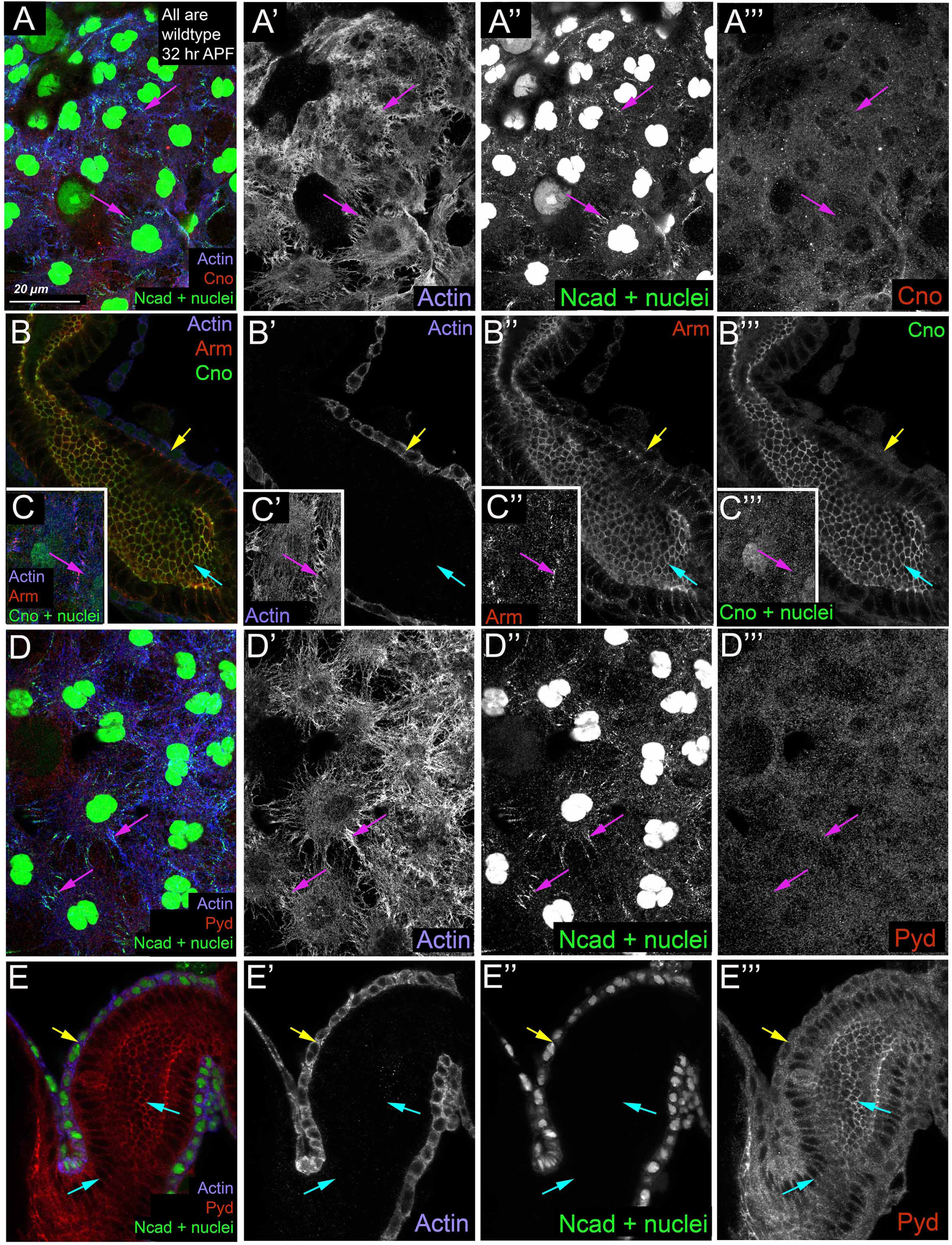
Ncad and Arm accumulate at migrating TNM borders while Cno and Pyd do not. Fixed wildtype pupal tissue during myotube migration at 31-32 hours APF. Myotubes express *Lifeact::eGFP* (blue) and mCherry^nls^ (revealing nuclei; green), both under the control of *mef2-Gal4*. Antigens used are indicated. **(A, C, D)** Front of the migrating myotube sheet. Magenta arrows denote Ncad-based myotube cell-cell junctions on the testis body. **(B, E)** Medial sections of the seminal vesicle. Yellow arrows denote myotubes surrounding the seminal vesicle. Cyan arrows denote cell-cell junctions of the seminal vesicle. **(A-C)** While Ncad and Arm localize to cell junctions between myotubes (A,C magenta arrows), Cno is not detected there. However, both Arm and Cno localize to cell junctions in the seminal vesicle (B, cyan arrows). **(D, E)** While Ncad localizes to cell junctions between myotubes (D, magenta arrows), Pyd is not detected there. However, Pyd localizes to cell junctions in the seminal vesicle (E, cyan arrow).

### Identifying and employing reagents for gene knockdown

We next identified shRNA reagents targeting the genes of interest. In the case of genes where null alleles are adult viable, we also obtained those alleles. In selecting shRNA reagents from the genome wide collections available from the Bloomington and Vienna Stock Centers, we prioritized those thought to work well in somatic tissues (Dietzl *et al*., 2007; Ni *et al*., 2011). When possible, we selected shRNAs with known phenotypes in somatic tissues (as described in the “Phenotypes” tab FlyBase (Ozturk-Colak *et al*., 2024); Suppl Table 2 right column). For a subset of the lines targeting genes with well-known effects in the embryonic epithelium, we further verified knockdown by looking for embryonic lethality after expression using a maternal germline driver (Suppl Table 3).

We expressed each selected UAS-shRNA in a tissue specific way, using the muscle-specific *mef2*-GAL4 driver (Duffy, 2002; Bischoff *et al*., 2025b). A few lines led to embryonic or larval lethality, precluding analysis. Others led to pharate adult lethality but at a stage where testis could be evaluated. Testis were dissected from seven to ten male progeny in a way in which their original chirality was not maintained, and stained with phalloidin, which binds to F-actin and outlines muscles. We then examined the testes under a dissecting microscope. At this level of resolution, the testis of many of our knockdowns were within the normal variability of wildtype (Suppl Fig 2A-J; Suppl Table 2)—for these a single testis was imaged via confocal microscopy. For shRNAs with phenotypes outside the wildtype range, we obtained ≥10 confocal images and tested the genotype at least twice. Using these images, each line was evaluated for both penetrance and severity (Suppl Table 2). Penetrance was by fraction with the phenotype: 0 = 0%, 1 = 10% - 25%, 2 = 26% - 50%, 3 = 51% - 75%, 4 = 76% - 100% with phenotype. As we observed in our previous screen (Bischoff *et al*., 2025b), many lines were incompletely penetrant. We do not know the causes of this, but one observation suggests it is not simply due to technical errors. We sometimes observed males in which one testis had a strong mutant phenotype and the other was wildtype (Suppl Fig 2K). Our knockdowns led to diverse defects in testis morphogenesis, including defects in distal muscle coverage, which may reflect issues with migration, gaps in muscle coverage along the proximal-distal axis, which may involve defects in adhesion, and defects in testis shaping, ranging from subtle differences in elongation to severe defects in shape. We thus scored severity as follows: 0 = no phenotype, 1 = mild distal testis phenotype with no muscle coverage defects, 2 = mild distal phenotype + other phenotype (wavy surface, flipped tip), or mild muscle coverage defects, 3 = moderate to severe muscle coverage defects, or severe shaping defects with no gaps, 4 = severe shaping and muscle coverage/migratory defects.

### The core catenins work with Ncad in testis morphogenesis and Ecad plays a supporting role

The wildtype testis is elongated along the proximal-distal axis, has a characteristic and spiral shape, and is completely covered in circumferential muscles (Fig 3A). *Ncad* knockdown has a characteristic phenotype, with gaps in muscle coverage all along the proximal-distal axis (Rothenbusch-Fender *et al*., 2017; Bischoff *et al*., 2025b)—we replicated this, with similar phenotypes generated by four shRNAs (Fig 1D-G, 3B). The cytoplasmic tails of all classical cadherins bind a conserved set of “catenins” that are critical for their function. These include Arm and α-catenin. Loss of Ecad (Tepass *et al*., 1996; Uemura *et al*., 1996)), Arm (Cox *et al*., 1996), or α-catenin (Sarpal *et al*., 2012) all lead to severe defects in embryonic morphogenesis. Arm also facilitates Ncad function in the embryonic nervous system (Iwai *et al*., 1997; Loureiro and Peifer, 1998). We thus hypothesized that the core catenins would be key for Ncad function in myotube migration. Previous work revealed a role for Arm there (Rothenbusch-Fender *et al*., 2017).

**Fig 3.**
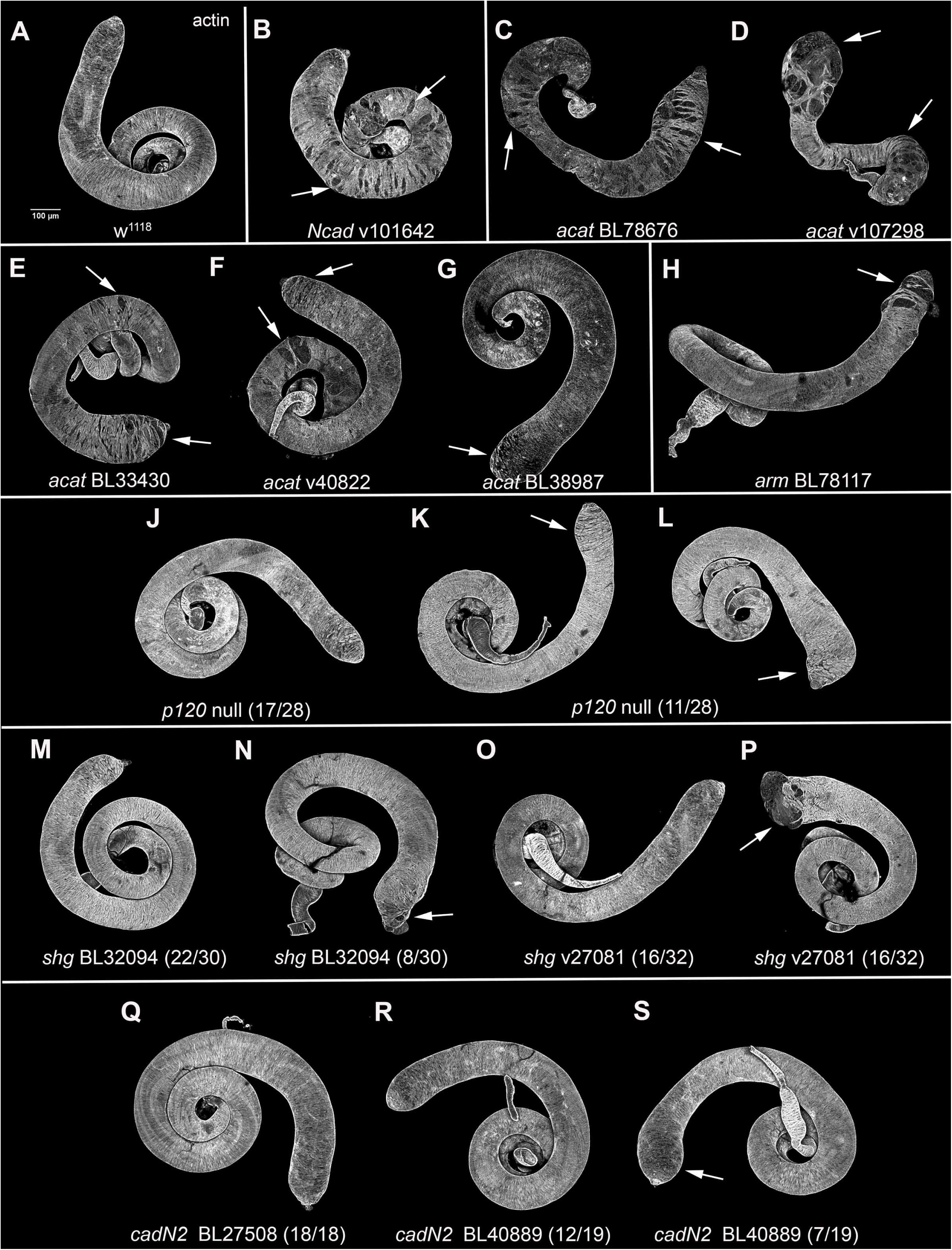
The core catenins and classical cadherin family proteins contribute to proper testis morphogenesis. **(A-S)** Testis from wildtype adults or adults expressing indicated shRNAs or null alleles. All are stained with fluorescently-labeled phalloidin to reveal F-actin, highlighting muscles. shRNA lines or genotypes are indicated. Arrows denote defects in adult testis morphogenesis. More detailed descriptions of phenotypes are in the text. (A) Wildtype (*w^1118^*), showing normal muscle coverage and shape. (B) Ncad knockdown, with gaps all along the proximal-distal axis. (C-G) Five a*-catenin* shRNAs had muscle coverage defects. (H) *arm* shRNA with distal muscle coverage defects. (J) 17/28 p120 null allele testis were wildtype in phenotype (K-L) 11/28 p120 null allele testis had distal testis broadening. (M) 22/30 *shg* shRNA BL32094 testes were wildtype. (N) 8/30 *shg* shRNA BL32094 testes had mild distal muscle coverage defects. (O) 16/32 *shg* shRNA V27081 testes were wildtype. (P) 16/32 *shg* shRNA B27081 testes had distal muscle coverage defects. (Q) 18/18 *cadN2* shRNA BL27508 testes were wildtype. (R) 12/19 *cadN2* shRNA BL40889 testes were wildtype. (S) 7/19 *cadN2* shRNA BL40889 testes had mild distal testis broadening. C-G are included in our Class 2 (cell adhesion defects). H and M-P are included in our Class 3 (distal muscle coverage defects). J-L and Q-S are included in our Class 4 (testis shaping defects).

We tested five shRNA lines targeting α-catenin. Four resembled Ncad knockdown, with gaps in muscle coverage along the proximal-distal axis (Fig 3C-F), while one was less severe (Fig 3G). Arm plays dual roles in cell adhesion and in Wnt signaling. Previous work revealed testis adhesion defects with the *arm* shRNA BL31304 (Rothenbusch-Fender *et al*., 2017)). We tested additional shRNAs. Two led to lethality before pupation (Suppl Table 2), potentially due to essential roles in Wnt signaling. Another shRNA gave defects in distal muscle coverage (Fig 3H) while one was wildtype (Suppl Table 2). We suspect that the dual roles of Arm mean that just the right amount of knockdown is required to both avoid early lethality and still achieve enough reduction to affect function. However, with this caveat, these data are consistent with essential roles for the Ncad/catenin complex in myotube migration.

Classical cadherins also bind members of the p120catenin family (Reynolds, 2007; Cadwell *et al*., 2016). Drosophila has a single family member. p120 proteins are critical for full classic cadherin function, stabilizing them at the plasma membrane and inhibiting endocytosis. *p120* mutant mice are embryonic lethal (Hernandez-Martinez *et al*., 2019). However, Drosophila *p120* null mutants are viable, though they have defects in embryonic development when the normal levels of E-cadherin are reduced (Myster *et al*., 2003), and have mild phenotypes in other times and tissues (Fox *et al*., 2005; Bulgakova and Brown, 2016; Liang *et al*., 2017; Iyer *et al*., 2019). We examined *p120* null homozygotes. Most testes were within the wildtype range (Fig 3J; 17/28), while others had mild to moderate distal enlargement (Fig 3K,L; 11/28 testes). Thus, as in embryonic development, the *p120* phenotype was milder than that of the classical cadherin.

Our previous work suggested Ncad was the primary classical cadherin in action in testis morphogenesis. *Ecad* (*shg)* mRNA was also present, but Ecad protein accumulated at low enough levels that we did not detect it at junctions during migration (Bischoff *et al*., 2025a). To explore whether Ecad might play a more modest role in testis morphogenesis, we tested three lines targeting Ecad (*shg)*. Two shRNAs, both previously validated (Suppl Table 2), had similar mild to moderate defects in distal muscle coverage and shaping (Fig 3M-P)—one was more penetrant than the other (V27081 16/32 testes; BL32904 8/30 testes). The other gRNA line was wildtype in phenotype (v342699; Suppl Table 2). This suggests that Ecad plays a modest role in migration or maintaining adhesion distally, unlike the broader role of Ncad in adhesion all along the proximal-distal access. Finally, we examined the second Ncad relative, CadN2. Our RNAseq data suggests it may not be expressed in this tissue (Table 1). We tested two shRNAs targeting CadN2. One was wildtype (Fig 3Q; BL27508; 18/18 testes), while the other had a very mild and moderately penetrant distal testis broadening (Fig 3R,S, BL40889 (7/19 testes). Given this and the low level of expression, we think it is unlikely CadN2 plays a significant role in testis morphogenesis.

### Several key regulators of embryonic morphogenesis have at most modest roles in the testis

Three of the proteins on our list, Cno (the Adadin homolog, bait in our proximity proteomics screen), Pyd (homolog of ZO-1 and the top hit in our proximity proteomics screen), and Baz (homolog of Par3) localize to embryonic AJs and play important roles in embryonic morphogenesis (Müller and Wieschaus, 1996; Sawyer *et al*., 2009; Choi *et al*., 2011). We thus were particularly curious to test whether they are also important for myotube migration and testis morphogenesis. We tested four shRNA lines targeting Cno. One strongly reduces Cno function when expressed in embryos (BL33367; (Manning *et al*., 2019) and in pupal eyes (Walther *et al*., 2018), and a second was validated in the brain (v7769; (Perez-Gomez *et al*., 2013). Both of these shRNAs, along with BL38194, had no effect on testis morphogenesis when expressed using *mef2*-GAL4 (Fig 4A vs C-E; n=8-11 images per genotype; Suppl Table 2). However, a fourth shRNA (v102686) had moderately penetrant mild distal defects (Fig 4F; 6/11 testes). To further explore possible roles for Cno in the testis, we obtained a strong hypomorphic mutant, *cno^mis^*, which is viable but has strong defects in adult eyes, wings and bristles (Miyamoto *et al*., 1995). We examined its phenotype both homozygous and when transheterozygous with a null *cno* allele, *cno^R2^.* Most testes in each genotype were within the wildtype range (e.g., *cno^mis^* homozygotes, 8/11; *cno^mis^/ cno^R2^,* Fig 4H, 11/15), but each had low penetrance mild distal testis defects (Fig 4G, I). Together, these results suggest that despite Cno’s important roles in many epithelial tissues, it does not play an essential role in Ncad-mediated myotube adhesion. It may play a modest role.

**Fig 4.**
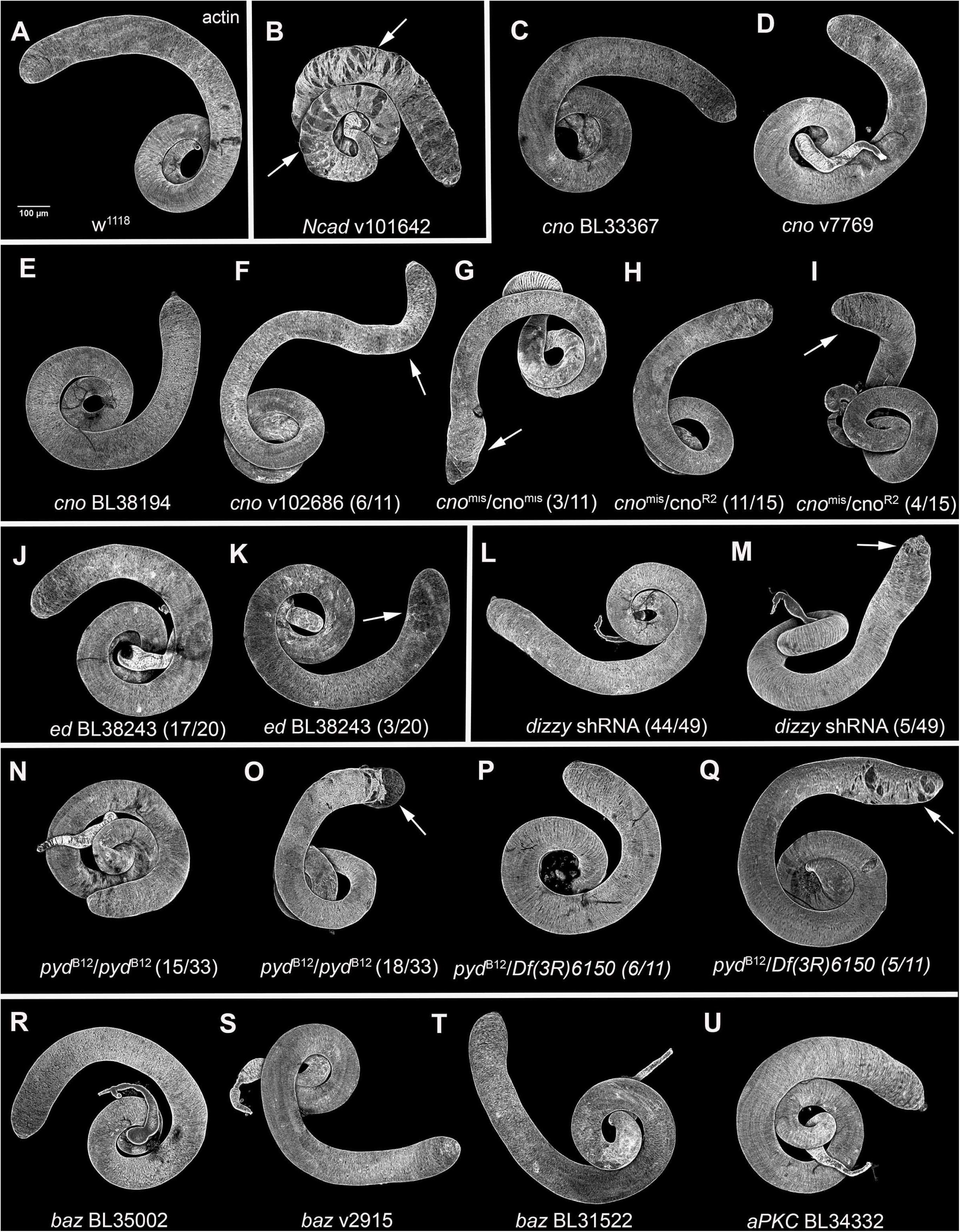
Cno, Pyd, Ed, Dzy, Baz, and aPKC play at most modest roles in testis morphogenesis. Testis from wildtype or adults expressing indicated shRNAs or null alleles, all stained with phalloidin to reveal F-actin. shRNA lines or genotypes are indicated. Arrows denote defects in adult testis morphogenesis. More detailed descriptions of specific phenotypes are in the text. (A) Wildtype (*w^1118^*), showing normal muscle coverage and shape. (B) Ncad knockdown. (C-E) Three shRNAs targeting Cno were wildtype in phenotype. (F) *cno* shRNA V102686 had a partially penetrant flipped distal testis (6/11). (G, H) 3/11 *cno^mis^/cno^mis^* testis had distal testis defects, while 11/15 were wildtype. (I) 4/15 *cno^mis^/cno^R2^* testis had distal testis broadening, while 11/15 were wildtype. (J, K) 17/20 *ed* shRNA testis were wildtype, while 3/30 had mild distal testis broadening. (L, M) 44/49 *dzy* shRNA were wildtype, while 5/49 had mild distal testis defects. (N, O) 15/33 pyd^B12^/pyd^B12^ testis were wildtype while 18/33 had distal muscle coverage defects. (P, Q) 6/11 *pyd^B12^/Df(3R)6150* testis were wildtype, while 5/11 had distal muscle coverage defects. (R-T) In cases where we examined mutants, all tissues are affected and we cannot rule out roles in tissues other than myotubes. Three shRNAs targeting Baz were wildtype in phenotype. (U) *aPKC* shRNA with a wildtype phenotype. C-K are included in our Class 4 (testis shaping defects). L-Q are included in our Class 3 (distal muscle coverage defects).

In parallel, we examined roles of the known Cno/Afadin binding partner and nectin relative Echnoid (Ed; (Wei *et al*., 2005), and of a known upstream activator of Cno, the Rap-guanine nucleotide exchange factor (GEF) PDZ-GEF/Dizzy (Bonello *et al*., 2018; Perez-Vale *et al*., 2023). Two shRNA lines targeting Echinoid were respectively wildtype (Suppl Table 2) or had very mild, low penetrance defects (Fig 4J, K; 17/20 wildtype; (Bischoff *et al*., 2025b). We used a validated shRNA to knockdown Dizzy; to enhance potential effects these animals were also heterozygous for a *dzy* null allele. Once again, most testes were wildtype (44/49; Fig 4L), with ∼10% having mild distal defects (Fig 4M). Thus, the results with Ed and Dizzy knockdown are consistent with what we observed with Cno, suggesting at most modest roles. The small GTPase Rap1 binds Cno and activates it (Boettner *et al*., 2003). In contrast, Rap1 plays an important role in testis morphogenesis (Bischoff *et al*., 2025a). However, Rap1 has multiple downstream effectors and in embryos its loss leads to more severe defects than loss of Cno (Perez-Vale *et al*., 2023), potentially due to defects in integrin-based adhesion (Ellis *et al*., 2013).

To test Pyd’s role, we used a protein null deletion allele, *pyd^B12^*, which is adult viable (Choi *et al*., 2011). We examined *pyd^B12^*/*pyd^B12^* homozygous mutants and *pyd^B12^*transheterozygous to two deletions removing *pyd* (*Df(3R)Exel6150* and *Df(3R)ED5296). pyd^B12^/Df(3R)ED5296* males had strong defects in the paragonia (the paired male accessory sex glands), with apparent fusion to the proximal testis (Suppl Fig 2M, N magenta arrows), making it difficult to fully assess the testis phenotype in this genotype. In *pyd^B12^* homozygotes many testes were wildtype (Fig 4N; 15/33 testes) while others had defects in distal muscle coverage or arrangement (Fig 4O; 18/33 testes). *pyd^B12^/ Df(3R)Exel6150* animals also had partially penetrant defects in distal muscle coverage (Fig 4P, Q; 5/11 testes). Since *pyd^B12^* is a null mutant, this reveals that Pyd’s role in testis morphogenesis is limited. It facilitates distal testis shaping but does not have an apparent role in adhesion more proximally. It is important to note that the *pyd^B12^* deletion also removes one other predicted gene CG8379, so even the phenotypes we observed could result from its deletion.

Finally, we tested potential roles for the apical polarity regulator Baz (fly Par3). We used three shRNA lines, all with validated function in embryos (BL35002; Suppl Table 3) or somatically (v2915 and BL31522; Suppl Table 2). None of these shRNAs led to defects in testis migration or morphogenesis when expressed using *mef2*-GAL4 (Fig 4R-T; n=11-23 images per genotype; Suppl Table 2), consistent with the idea that Baz does not play an important role in this mesenchymal tissue. Given this, we also tested three shRNAs targeting aPKC, another key apical polarity regulator. Once again, two have been verified in multiple contexts (Suppl Tables 2 and 3) but all three were wildtype in phenotype (Fig 4U; Suppl Table 2; n=1-10 images per genotype). Together, these data reveal that several key regulators of epithelial cell morphogenesis in embryos play at most modest roles on myotube mesenchymal migration and morphogenesis.

### Defining the subset of junctional proteins that function in myotube mesenchymal migration

Having examined the core catenins and three key regulators of epithelial morphogenesis, we then examined the roles of the 55 remaining candidates from our proteomics list using shRNAs/gRNAs or null alleles where available. We tested 116 shRNA/gRNA lines and 10 mutant/null allele lines. It’s important to note in advance that a negative result with any shRNA does not rule out a role for that protein in morphogenesis—while we attempted to select validated shRNAs, this does not guarantee efficient knockdown in our tissue and stage. In those cases in which only a single shRNA had a phenotype, further work is needed to ensure this is not an off-target effect. From our 58 candidates, we identified strong to moderat morphological phenotypes in 22 genes.. These varied widely, from strong disruption of testis morphogenesis to milder defects in muscle coverage or testis shaping. We broadly categorized adult testis phenotypes into the following categories (Table 2): severe loss of muscle coverage (Class 1), major testis cell adhesion defects (Class 2), distal muscle coverage defects (Class 3), and testis shaping defects (Class 4).

### Par-1, Dlg5, Rap1, and RN-tre knockdown each severely alter adult testis morphogenesis

Among all of our knockdowns, perhaps the most dramatic phenotype was that caused by knockdown of the Ser/Thr kinase and basolateral polarity regulator Par1 (Table 2; (Wu and Griffin, 2017). Par-1 knockdown led to extremely severe and highly penetrant loss of muscle coverage, with the testes failing to elongate and begin the spiralization process. We observed this strong, highly penetrant phenotype in two of the three shRNA/gRNAs targeting Par-1 (Fig5A vs. B,C: BL32410 (36/40 testes), V341094 (17/19 testes). This was a phenotype we had only observed once before, after knockdown of Drosophila cytoplasmic myosin (Rothenbusch-Fender *et al*., 2017; Bischoff *et al*., 2021).

The other three drastic phenotypes involve severe shaping and/or severe cell adhesion defects. Discs Large 5 (Dlg5) is a MAGUK protein with known roles in epithelial polarity and border cell migration. Dlg5 knockdown with one shRNA had a highly penetrant phenotype with severe gaps all along the proximal-distal axis (Fig 5D; BL30926; 19/20 testes), similar to but more severe than those caused by to Ncad knockdown. Defects were even more severe with a second shRNA (BL61334). This caused extensive loss of muscle coverage like that we saw with Par-1 knockdown, and led to testis shaping defects that varied from an oval testis (Fig 5F; 6/17 testes) to a round testis (Fig 5E; 11/17 testes; Table 2), similar to what we observed when knocking down the RhoGEF PIX (Bischoff *et al*., 2025b).

**Fig 5.**
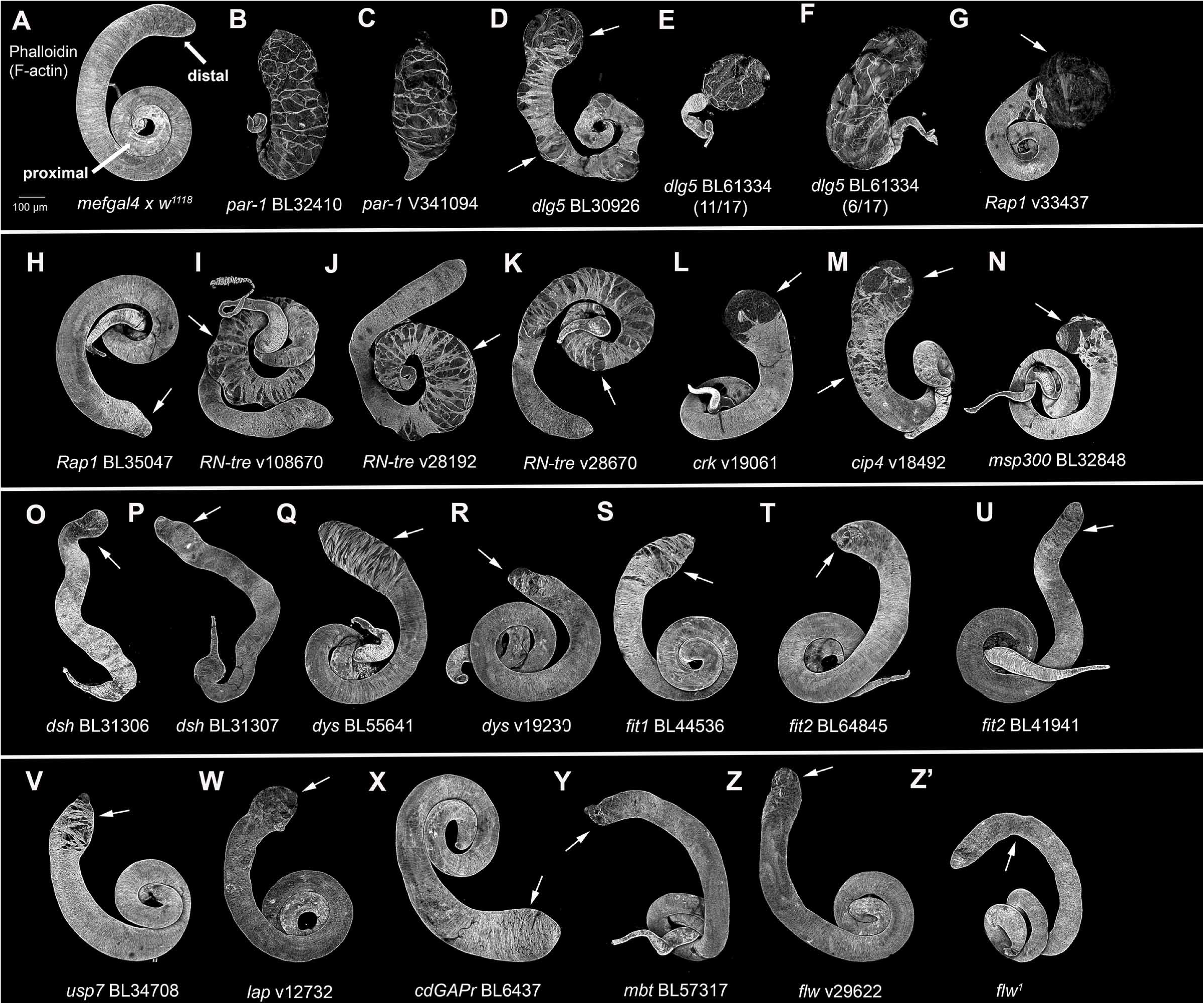
Some gene knockdowns have major impacts on muscle coverage, testis shaping or cell adhesion in the adult testis. Testis from wildtype or adults expressing indicated shRNAs or mutant alleles. All are stained with phalloidin to reveal F-actin, highlighting muscles. shRNA lines or genotypes are indicated. Arrows denote defects in adult testis morphogenesis. More detailed descriptions of specific phenotypes are in the text. (A) Wildtype (*mef2-Gal4* crossed with *w^1118^*). Arrows indicate proximal and distal testis. (B, C) Two *Par-1* shRNAs with extreme loss of muscle coverage and failure to spiralize. (D-F) Two *dlg5* shRNAs with extreme loss of muscle coverage and shaping defects. (G, H) Two *Rap1* shRNAs. One has severe distal muscle coverage defects and the other has mild distal defects. (I-K) Three RN-tre shRNAs. All lead to proximal testis cell adhesion defects. (L-N) *crk, cip4,* and *msp300* shRNAs all lead to severe distal muscle coverage defects. (O, P) Two *dsh* shRNAs lead to shaping defects and failure to spiralize. (Q, R) Two *dys* shRNAs with moderate defects in distal muscle coverage. (S) Fit1 knockdown leads to moderate defects in distal muscle coverage. (T, U) Two *fit2* shRNAs. One has mild distal muscle coverage defects and the other has mild shaping defects. (V) Usp7 knockdown leads to moderate defects in distal muscle coverage. (W) Lap knockdown leads to mild defects in distal muscle coverage. (X) cdGAPr knockdown leads to a broadened distal testis. (Y) Mbt knockdown leads to mild defects in distal muscle coverage. (Z-Z’) One *flw* shRNA leads to mild defects in distal muscle coverage while *flw^1^* homozygotes have a mild wavy testis surface. B-F are included in our Class 1 (severe loss of muscle coverage). I-K are included in our Class 2 (cell adhesion defects). G-H, L-N, and Q-Aa are included in our Class 3 (distal muscle coverage defects). O-P are included in our Class 4 (testis shaping defects).

Rap1 is a small GTPase that regulates cell adhesion and junction formation. Rap1 knockdown led to severe muscle coverage defects (Fig 5G, V33437; 18/20 testes); this Rap1 line and a small number of other lines noted below overlapped between this screen and our earlier work (Bischoff *et al*., 2025a; Bischoff *et al*., 2025b). A second Rap1 shRNA had a milder, less penetrant phenotype with the distal testes constricted (Fig 5H; BL35047 (7/19 testes)).

Knockdown of the Rab5 GAP RN-tre, which has known roles in regulating junctional protein trafficking, had a quite different effect. RN-tre knockdown led to highly penetrant cell adhesion defects (Fig 5I-K; Table 2). However, in contrast to Ncad knockdown, which leads to gaps all along the proximal-distal axis (Fig 1D-G; (Bischoff *et al*., 2025a), the gaps seen after RN-tre knockdown were concentrated entirely in the proximal half of the testis (Fig 5 I-K, arrows) while the distal testis was unaffected, Similar penetrant defects were seen with three of the four lines we tested (V108670 (19/21 testes); V28192 (16/18 testes; BL28670 (11/18 testes). This proximally restricted adhesion defect was not observed in our previous work.

### shRNAs targeting multiple candidates alter distal muscle coverage

Another group of knockdowns led to phenotypes involving myotubes that failed to migrate to and/or adhere to one another to cover the distal testes in muscle (Table 2). These phenotypes varied in severity. Knockdown of several genes led to severe distal defects, with no muscle coverage on the distal testis. These included knockdown of the adaptor protein Crk (Fig 5L; V19061 (12/19 testes), the F-bar domain protein Cip4 (Fig 5M, V18492 (16/20 testes); also examined in (Bischoff *et al*., 2025b)), and of MSP300, a KASH domain Nesprin family member that regulates actin-dependent nuclear anchorage (Fig 5N; BL32848 (5/20 testes)). Disheveled (Dsh) plays roles in Wnt signaling and planar cell polarity, and its knockdown had a different phenotype. Two shRNA lines targeting Dsh led to highly penetrant shaping and muscle alignment defects (Fig 5O,P; BL31306 (8/8 testes), BL31307 (8/8 testes)). These were similar to those observed when knocking down the Wnt ligand Wnt4 and its receptor Fz2 (Bischoff *et al*., 2025b).

Along with knockdowns leading to a severe loss in distal muscle coverage, we also observed more moderate phenotypes in which the distal testis was encompassed in muscle, but the muscle sheet at the tip had significant gaps. Three of these were proteins involved in linking the extracellular matrix to the cytoskeleton: Dystrophin (Fig 5Q,R; targeted by BL55641 (13/20 testes) and V19230 (2/18 testes)), and Fermitin1 (Fit1) and Fermitin 2 (Fit 2), Kindlin family members that play important roles in integrin-based signaling (Fit1; Fig 5S; V44536 (12/20 testes); Fit2; Fig 5T; BL64845 (5/20 testes). A second *fit2* shRNA had milder defects (Fig 5U; BL41941 (4/19 testes)). Similar moderate distal muscle coverage defects were seen with three other proteins of diverse functions: the ubiquitin-specific protease USP7 (Fig 5V; BL34708 (4/19 testes)), the clathrin adaptor protein Like-AP180 (Lap; Fig 5W, V12732 (9/20 testes), and the Rho family GAP cdGAPr (Fig 5X; BL6437; 9/11 testes). shRNAs targeting two other genes gave milder defects in distal muscle coverage or shape. They targeted Mushroom bodies tiny (Mbt), the Drosophila homolog of Pak2 and Pak4 (Fig 5Y, BL57317 (7/15 testes)), and Flapwing (Flw), a protein-serine/threonine phosphatase (PP1β) that regulates nonmuscle myosin (Fig 5Z, B29622 (6/27 testes)). For Flapwing, we also tested a viable missense mutant, *flw^1^*, which had a mild wavy testis border and distal testis broadening (Fig 5Z’; 4/12 testes). Together, these data suggest the distal testis is most vulnerable to gene knockdown.

### Roles for Ena/VASP proteins and their regulator Abl

Our earlier work revealed roles for key actin regulators in migration, including the Arp2/3 complex and Formins (Bischoff *et al*., 2021). Enabled (Ena), another of our candidates, is also an actin regulator. As part of the Ena/VASP family, it helps drive actin polymerization. Two shRNAs targeting Ena led to a pointed distal testis with mild muscle coverage defects (Fig 6A vs. B,C: BL31582 (10/18 testes), V43056 (12/21 testes)). To confirm this was due to reduced Ena function, we expressed FP4mito, which recruits Ena to mitochondria and inactivates it (Gates *et al*., 2007). FP4mito expression also led to a pointed testis with mild muscle coverage defects (Fig, 6D; 10/19 testes). We looked further into this pathway by exploring a role of the non-receptor tyrosine kinase Abl, which negatively regulates Ena (Grevengoed *et al*., 2001). We tested two *abl* shRNAs and both were wild-type (Suppl Table 2). However, we also expressed a kinase-dead form of Abl (Wills *et al*., 1999) using *mef2*-GAL4. This led to highly penetrant, severe defects in the distal testis migration (Fig 6E; BL8566 (17/17 testes). These new data extend our understanding of roles of regulated actin polymerization in testis morphogenesis.

**Fig 6.**
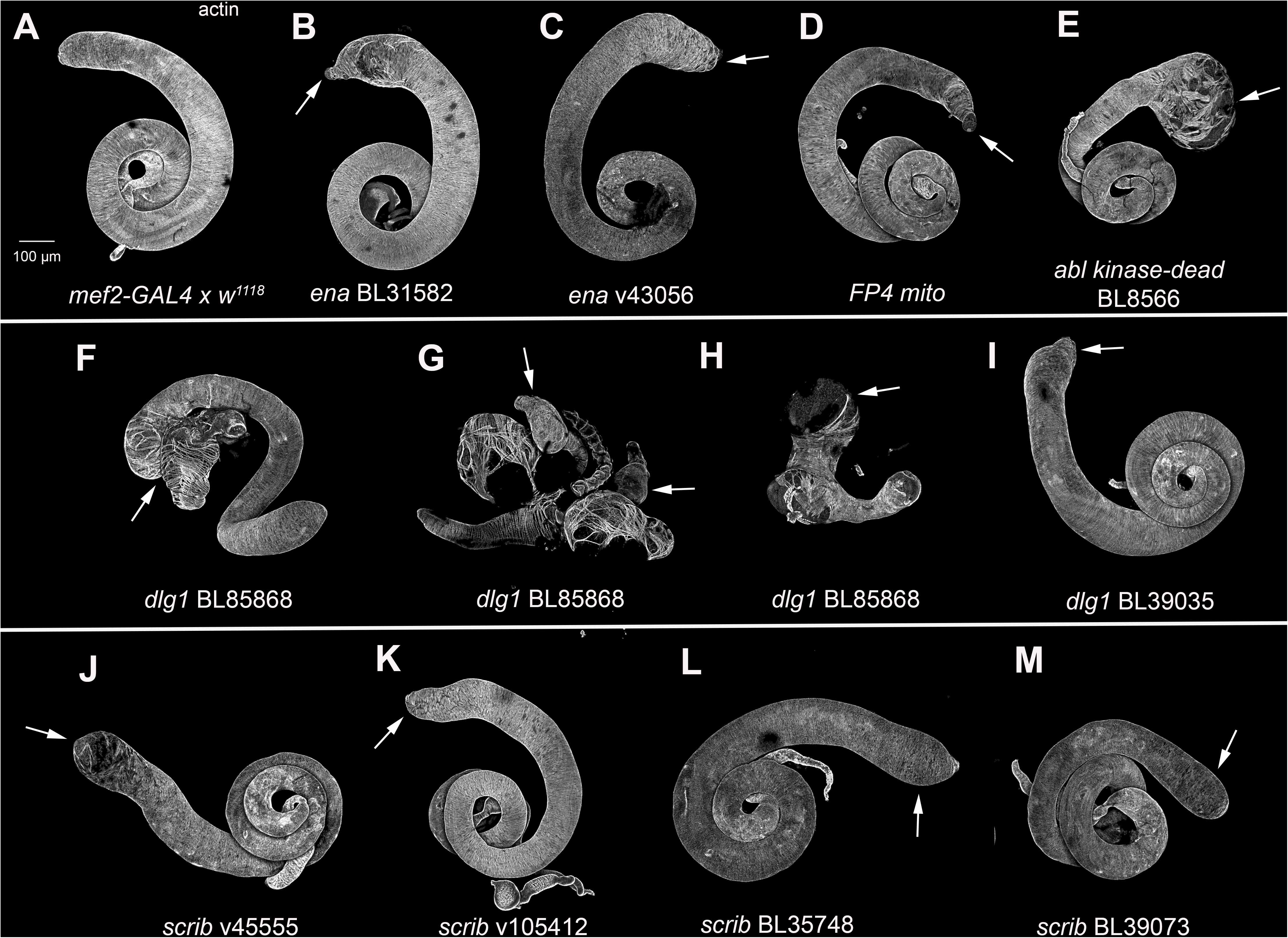
The Ena/VASP family and some basolateral polarity proteins play roles in testis morphogenesis. Testis from wildtype or adults expressing indicated shRNAs or mutant constructs. All are stained with phalloidin to reveal F-actin, highlighting muscles. shRNA lines or genotypes are indicated. Arrows denote defects in adult testis morphogenesis. More detailed descriptions of specific phenotypes are in the text. (A) Wildtype. (B-C) Two *ena* shRNAs both led to a pointed distal testis and mild muscle coverage defects. (D) Expressing FP4mito to sequester Ena also leads to a pointed distal testis and mild muscle coverage defects. (E) Overexpressing an Abl kinase dead mutant leads to severe distal muscle coverage defects. (F-H) *dlg1* shRNA BL85868 leads to fusion of the testis to paragonia (F, arrow) and severe distal shaping and muscle coverage defects in the testis (G, H arrows). (I) A second *dlg1* shRNA leads to distal testis broadening. (J-M) Four *scrib* shRNAs all lead to mild distal testis muscle coverage or shaping defects. B-E and J-M are included in our Class 3 (distal muscle coverage defects). F-I are included in our Class 4 (testis shaping defects).

### Two additional basolateral polarity proteins regulate testis morphogenesis

Drosophila epithelial basolateral polarity is regulated by the scaffolding proteins Dlg, Scrib and Lgl. (Schmidt and Peifer, 2020). Par1, described above, acts in parallel to phosphorylate and thus expel apical polarity proteins from the basolateral domain (Wu and Griffin, 2017). We tested two shRNAs targeting *lgl*, one with a validated phenotype in follicle cells, but neither affected testis morphogenesis (Suppl Table 2). In contrast, two of the five *dlg* shRNAs we tested had phenotypes. The first had a complex and very strong effect, in which the adult testis appeared to be fused to the paragonia (Fig 6F, arrow). While this made phenotypic analysis of the testis itself more challenging, they included examples where the testis failed to elongate and/or muscle coverage was severely affected (Fig 6G, H, arrows; BL85868 (17/23 testes)). A second *dlg1* shRNA had moderate distal testis shaping defects (Fig 6I: BL39035;11/19 testes). We also tested five *scrib* shRNAs, of which four had phenotypes. Two shRNAs led to mild defects in distal muscle coverage or distal muscle alignment (Fig 6J, K; V45555 (19/45 testes), V105412 (17/24 testes)). The other two lines showed a mild broadening of the distal testis (Fig 6L, M; BL35748 (8/33 testes), BL39073 (8/24 testes)). Together, these data suggest that the basolateral module plays a role in mesenchymal cell migration or testis shaping.

### Some shRNAs caused only mild testis shaping defects

A final set of knockdowns had mild defects in the shape of the distal testis but left the musculature fully intact (Suppl Fig 3B-U). Many of these defects were modest and some were low penetrance, but all were out of the range of sporadic developmental defects we see in wildtype (Suppl Fig 3A-J). We describe each in more detail in the legend for Suppl Fig. 3 In some cases we observed phenotypes with more than one shRNA line: these included knockdown of Shroom (Shrm; Suppl Fig 3B-E), a regulator of myosin contractility, and Zyxin (Suppl Fig 3F, G) and Ajuba (Jub; Suppl Fig 3H, I), LIM domain proteins with roles in both cell-cell and cell-matrix junctions. In the case of Outspread (Osp), the homolog of MPRIP (Myosin Phosphatase Rho-Interacting Protein), we found phenotypes with both an shRNA line and a viable hypomorphic allele, *osp^1^*(Suppl Fig 3S,T) Further work will be needed for these to determine whether these mild defects are due to modest roles, shRNA lines that only lead to partial knockdown, or off-target effects. One other shRNA line was puzzling, as its phenotype was quite variable (Suppl Fig 3V, W). This line targeted CG7600, the fly homolog of FAM91A1, a component of the WDR11 complex that regulates Golgi trafficking. In multiple but not all replicates, we found a subset of animals that died as pharate adults with moderate to strong phenotypes (Suppl Fig 3V, W), but another subset that eclosed and were wildtype or had mild distal broadening. This may be another reflection of the variability in penetrance we observed with other lines.

### Null alleles allowed us to rule out roles for the ASPP/Rassf8/Magi complex

Our shRNA approach cannot rule out a role for a protein, without independent confirmation that the shRNA eliminated protein expression. However, for some of our candidates, molecular null alleles existed, allowing us to provide a definitive test of function. One set of candidates of particular interest to us were the scaffolding proteins in the ASPP/RASSF8/Magi complex. They bind to one another and play important roles in the complex cell shape changes that assemble the stereotyped ommatidia in the fly eye (Langton *et al*., 2009; Zaessinger *et al*., 2015). This tissue utilizes both Ecad and Ncad (Hayashi and Carthew, 2004), and Cno also plays an important role there (Matsuo *et al*., 1997). To test if these epithelial scaffolding proteins play a role in migrating mesenchymal tissue, we examined the adult testes from validated null mutants. Flies homozygous for the null alelle *Magi^ex214^* (Zaessinger *et al*., 2015) had wildtype testes (Fig 7A vs B; 15/15 testes), as did two shRNA lines targeting Magi (Suppl Table 2). Males homozygous for the null allele *RASSF8^6^* also had wildtype testes (Fig 7C; 8/8 testes), as did animals expressing two shRNA targeting RASSF8 (Suppl Table 2). Most males homozygous for the null allele *ASPP^8^* (Langton *et al*., 2009) had wildtype testes (Fig 7D; 13/17) while a few had a mild, low penetrance wavy testis borders (Fig 7E; 4/17 testes). Likewise, males homozygous for *ASPP^d^*, which eliminates the ASPP longer isoform (Langton *et al*., 2009) were wildtype (Fig 7F; 14/14 testes), as were those expressing an shRNA or a gRNA targeting ASPP (Suppl Table 2). Thus, while these scaffolding proteins play important roles in the eye epithelium, they do not play an important role in testis morphogenesis.

**Fig 7.**
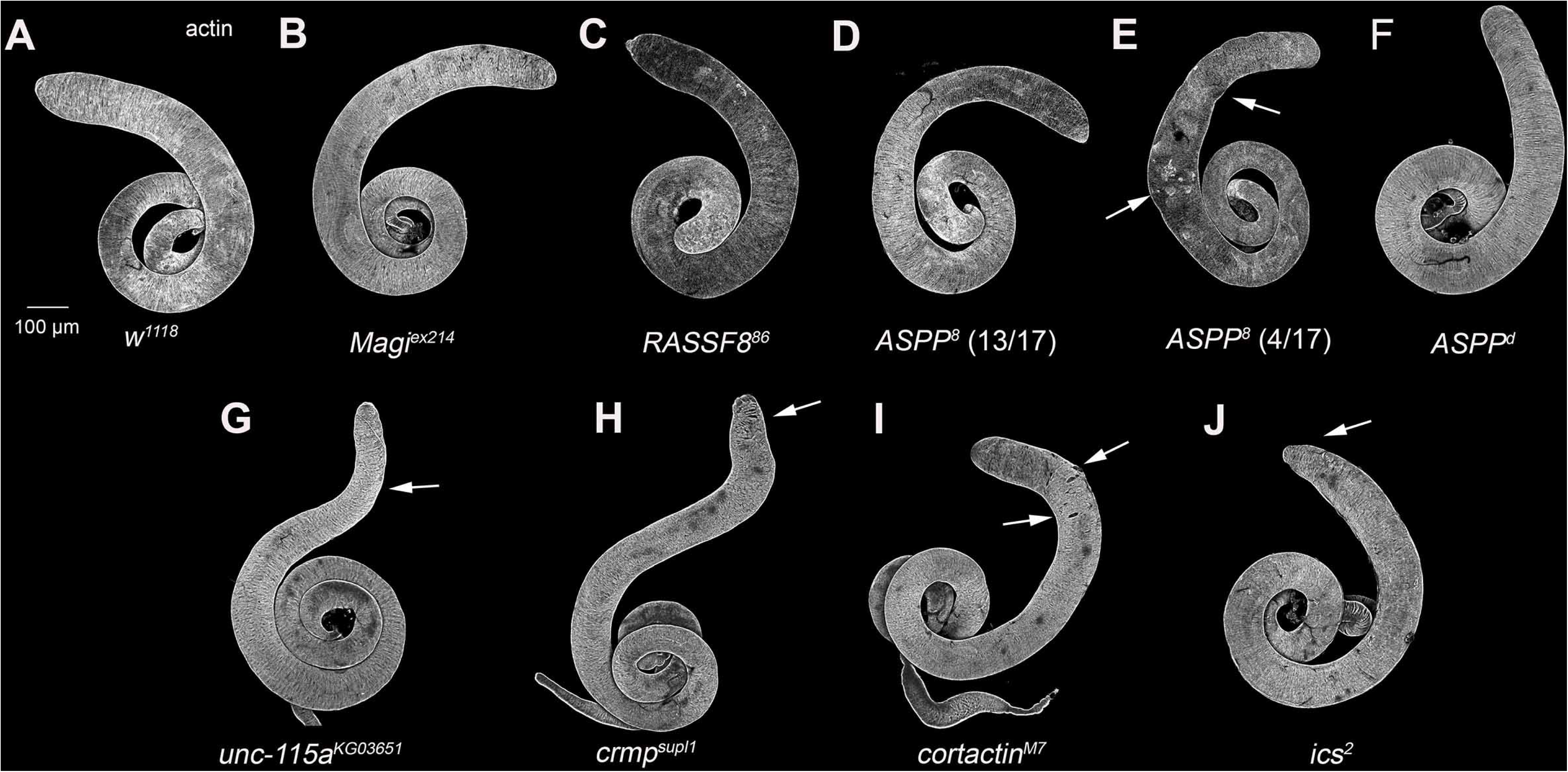
The use of null alleles allows us to rule out or limit roles of several of our candidates. Testis from wildtype adults and mutant adults. All are stained with phalloidin to reveal F-actin, highlighting muscles. Arrows denote defects in adult testis morphogenesis. More detailed descriptions of specific phenotypes are in the text. (A) Wildtype. (B-C) *Magi^ex214^* and *RASSF8^86^* mutants are wildtype in phenotype. (D, E) 13/17 *ASPP^8^* testes were wildtype, while 4/17 had wavy testis borders. (F) *ASPP^d^* mutant were wildtype. (G) *unc-115a^KG03651^*mutants had highly penetrant distal testis constriction, with a low penetrance flipped distal testis, (9/11 testes). (H) *crmp^supl1^* mutants had mild distal shaping defects (9/13 testes). (I) *cortactin^M7^* mutants had mild, low penetrance distal muscle coverage defects (3/11 testes) (J) *ics^2^*mutants had a mild constricted distal testis. (7/11 testes). D, E, G, H, and J are included in our Class 4 (testis shaping defects). I is included in our Class 3 (distal muscle coverage defects).

Several other candidate genes on our list had null alleles with mild phenotypes. These included Unc-115a, fly homolog of the ABLIM family of actin-binding proteins (Fig 7G; *unc-115a^KG03651^*(Garcia *et al*., 2007), Collapsin Response Mediator Protein (Crmp), an actin regulator whose mammalian relatives have roles in axon guidance (Fig 7H; *crmp^supl1^*(Morris *et al*., 2012), Cortactin, an Arp2/3 complex regulator (Fig. 7I; *cortactin^M7^*; (Somogyi and Rorth, 2004),, and Icarus, an adapter protein with a supporting role in integrin function (Fig 7J; ics^2^; (Green *et al*., 2018). Thus, these genes play relatively modest roles in testis morphogenesis but are not substantial players there. Taken together, our screen reveals roles for multiple AJ proteins in myotube collective migration and testis morphogenesis. These will serve as a resource for the community to further explore underlying mechanisms.

### While Par-1 and RN-tre knockdown both dramatically alter adult testis architecture, only Par-1 knockdown has effects during migration and these parallel effects of myosin knockdown

To illustrate how our screen will help illuminate the roles of AJ proteins in collective cell migration and testis morphogenesis, we explored two of our strongest hits in more detail: RN-tre and Par-1. Both had phenotypes with more than one shRNA/gRNA, and each had a phenotype not seen in other hits in our screen. RN-tre knockdown had a novel phenotype: gaps in the muscles restricted to the proximal testis (Fig 8A vs B). In this it was unlike many of our hits that preferentially affected the distal testis, and unlike Ncad knockdown that led to gaps all along the proximal-distal axis. Par-1 knockdown led to complete failure to enclose the testis in muscle and thus failure to re-shape it into the elongated spiral seen in wildtype (Fig 8C). This was a phenotype we had only seen once before—after knockdown of either the cytoplasmic myosin heavy chain (*zipper*) or its regulatory light chain (*sqh*) (Bischoff *et al*., 2021) (Bischoff *et al*., 2025a)

**Fig 8.**
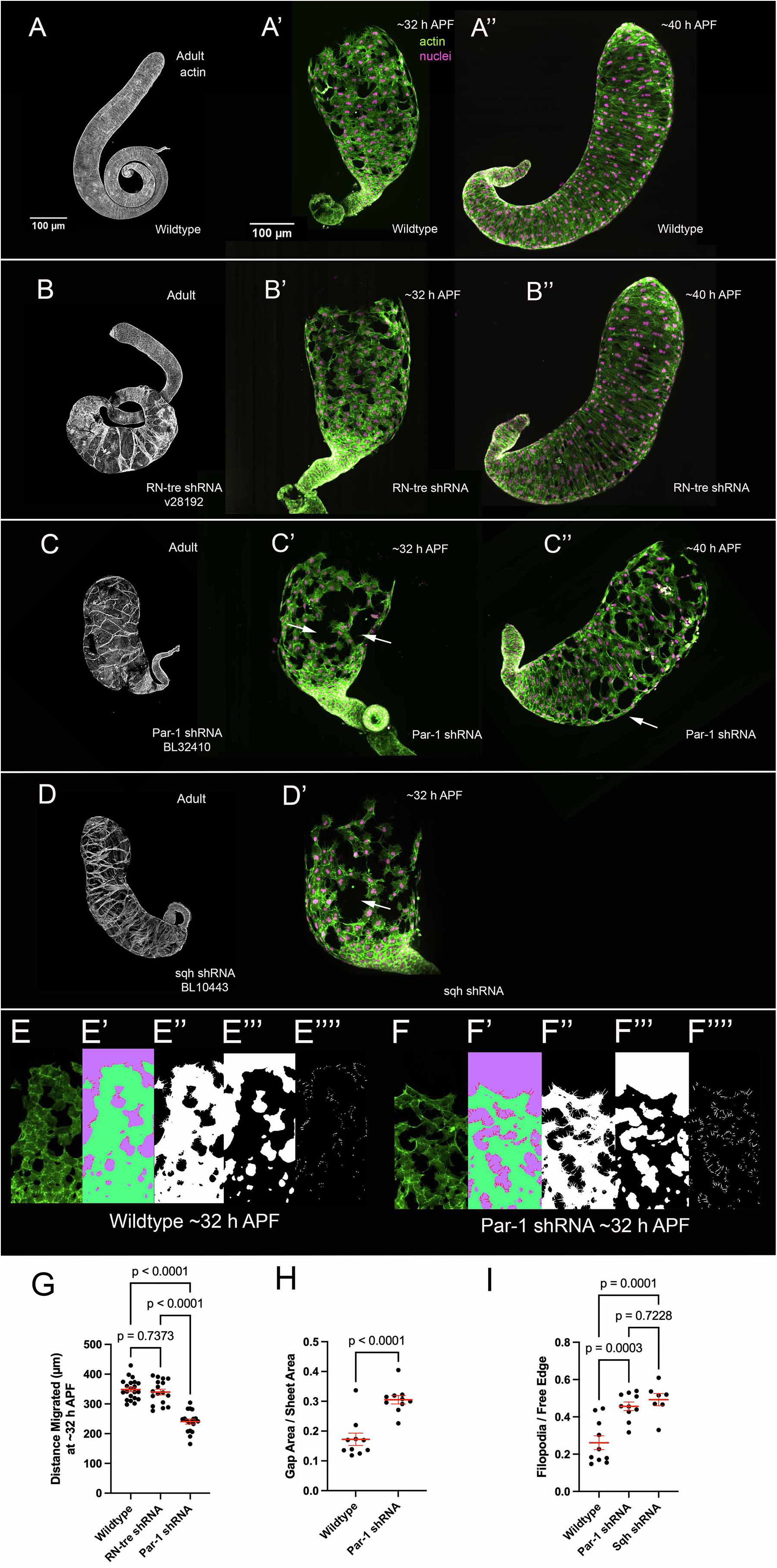
Live cell imaging reveals that Par-1 plays an important role in myotube migration. All adult testes were stained with fluorescently-labeled phalloidin to reveal F-actin, highlighting muscles. Stills from live cell imaging have myotubes expressing *Lifeact::eGFP* and mCherry^nls^ under the control of *mef2-Gal4,* with actin visualized in green and nuclei visualized in orange. **(A-A’’)** Wildtype adult testis (A), wildtype testis at 31-32 hours APF (A’), wildtype testis 40 hours APF (A’’). **(B-B’’)** RN-tre knockdown adult testis (B), RN-tre knockdown at 31-32 hours APF (B’), RN-tre knockdown at 40 hours APF (B’’). **(C-C’’)** Par-1 knockdown adult testis (C), Par-1 knockdown at 31-32 hours APF (C’), Par-1 knockdown at 40 hours APF (C’’). Arrows denote gaps in the TNM sheet during migration (31-32 APF) and elongation (40 APF). **(D-D’’)** Myosin regulatory light chain (Sqh) knockdown adult testis (B) and knockdown at 31-32 hours APF (D’) **(E-F’’’’)** Cropped images of migrating cells in 31-32 h APF myotubes and analysis. E, F. Cropped image. E’, F’ Labeled Image (Weka-Output) with three labels. red: filopodia, green: cell body, purple: background. E”, F’’ Binary image of TNM with filopodia (Sum of the red and green channels in E’). E’”, F’’’ Binary image of gaps (green channel from E’ or F’ inverted and smoothed with a morphological closing filter) E””, E’’’’ Binary image of filopodia (red channel from E’ or F’). **(G)** Average distance migrated in µm at 31-32 hours APF. **(H)** Ratio of gap area over total sheet area at 31-32 hours APF. **(I)** Ratio of filopodia area over free edge area at 31-32 hours APF. In each case the line represents the mean and error bars represent SEM, and all datapoints are indicated.

To determine the developmental timepoint at which these proteins first act, we imaged living explanted pupal testes beginning at 31-32 h APF, thus during migration (Bischoff and Bogdan, 2023). We used a stock expressing Lifeact-eGFP, visualizing F-actin and thus revealing cell shapes and protrusions, and mCherry with an added nuclear localization signal (NLS), highlighting nuclei. Both were under UAS-control and expressed specifically in myotubes using *mef2*-GAL4. Wildtype nascent myotubes migrate from the genital disc onto the testis, into the narrow space between the overlying pigment cells and the underlying germline cells (Fig 1A; Video 1). The cells migrate collectively (Fig 8A’), joined by Ncad-based junctions. Cells require attachment to the cohort to maintain directionality. Gaps form and close in the sheet by myosin-based contractility (Bischoff *et al*., 2021). When we examined testes after RN-tre knockdown, we observed no apparent changes in migration or gap closure relative to wildtype (Fig 8A’ vs B’; n=21 and 18 testes). In contrast, after Par-1 knockdown migration appeared delayed and gaps were more prominent (Fig 8A’ vs C’, arrows; Video 2; n=17 testes). To examine potential delays in migration, we quantified total distance migrated in stills taken at 31-32 h APF by measuring the distance from the bottom of the pupal testis (where it connects to the seminal vesicle) to the front of the migrating sheet (Fig 8G). By this time, wildtype myotubes had migrated 349±8 µm (n=21 testes). RN-tre knockdowns had similarly migrated 340±10 µm (n=17 testes). In contrast, Par-1 myotubes only had migrated 240±8 µm (n=18 testes). This suggests that myotube migration is delayed when Par-1 is knocked down, while RN-tre knockdown does not affect this. Intriguingly, nonmuscle myosin (sqh) knockdown also leads to delayed migration (Bischoff *et al*., 2021).

Next, we examined testis at 40 APF. At this stage in wildtype, migration to the distal end of the testis is completed, and myotubes are beginning to elongate to form the circumferential musculature (Fig 8A”; n=12 testes; (Bischoff and Bogdan, 2023), but the process of spiralization has only just begun. Intriguingly, RN-tre knockdown testis once again appeared similar to wildtype, with no apparent gaps (Fig 8B”; n=11 testes). To verify that this genotype had the observed adult phenotype, we let some animals progress to adulthood, and saw the characteristic proximal adhesion defects. This suggests that the defects seen after RN-Tre knockdown emerge during spiralization, which involves events in the proximal testis (Bischoff and Bogdan, 2023). In contrast, in animals with Par-1 knockdown the distal most testis was often uncovered, multiple gaps were present in the sheet and muscle elongation appeared delayed (Fig 8C”; n=21 testis).

As mentioned above, the adult phenotype of *par-1* shRNA knockdown resembled that of knockdown of non-muscle myosin (Fig 8D; (Bischoff *et al*., 2021; Bischoff *et al*., 2025a). To further define the nature of the defects after Par-1 knockdown and to compare them with defects we observed during migration after myosin, we used machine learning to segment stills of migrating myotubes at ∼32 hours APF (7-10 per genotype) and quantify gap areas well as other features. After training, the algorithm began with initial stills (Fig 8E, F). These were then segmented into regions corresponding to cell bodies, filopodia and regions unoccupied by cells (Fig 8E’, F’. E” vs E”’, F” vs F”’). The area occupied by cells thus resembles the sum of the cell body and filopodia regions (Fig 8E’’, F’’). We further extracted the filopodial area from the segmented picture (Fig 8E’’’’, F’’’’). To deduce the length and area of cell edges, we extracted the gap area from the segmented data and used a smoothing algorithm. Our first output assessed gap area as a fraction of the total area covered by the cell sheet. In wildtype, gaps covered 17±2% of the total area of the sheet (Fig 8H). As our visual assessment suggested, after Par-1 knockdown, gaps now increased to 30±1% of the total area of the sheet (Fig 8H). Myosin knockdown is also known to increase gaps relative to wildtype (Bischoff *et al*., 2021). In our previous analysis, we observed actomyosin cables that appear to play an important role in gap closure (Bischoff *et al*., 2021). Myosin knockdown leads to loss of these cables. Instead, the cells surrounding gaps have filopodia at all free edges (Bischoff *et al*., 2025a). As a proxy for the number of filopodia, we quantified filopodial area per free edge length after Par-1 knockdown. In wildtype, we observed 0.26±.0.04 µm²_filopodia_/µm_free edge_ while after Par-1 knockdown this increased to 0.46±0.02 µm²_filopodia_/µm_free edge_ (Fig 8I). Myosin knockdown had a similar increase in filopodia with 0.49±.0.03 µm²_filopodia_/µm_free edge_ (Fig 8I). Together, these data reveal multiple parallels between Par-1 and myosin knockdown. Intriguingly, Par-1 also plays a role in a different *Drosophila* collective cell migration event, border cell migration, where it is positively regulates myosin function by inhibiting myosin phosphatase (McDonald *et al*., 2008).

### Double knockdown suggests broader roles for RN-tre in adhesion

Both Ncad and RN-tre knockdown lead to small gaps in the sheet of circumferential muscles in adults. Given the known mechanistic role of Ncad in cell-cell adhesion, we suspect RN-tre might play a similar role. However, Ncad and RN-tre knockdown differ in the location of the gaps observed. Ncad knockdown leads to gaps all along the proximal-distal axis, while RN-tre knockdown only leads to gaps proximally, while none are seen distally. As one approach to further explore these potentially parallel roles, we compared single knockdown of each gene to knockdown of both simultaneously, controlling for UAS copy number.

Single knockdown of each gene with this expression control included replicated what we had seen during the screen. Ncad knockdown led to gaps all along the proximal distal axis (Fig 9A), while RN-tre knockdown only led to proximal and not distal gaps (Fig 9B, D, F, yellow vs cyan arrows). We quantified gaps by outlining them and assessing fraction of the testis area they occupied. We used three different RN-tre shRNAs with somewhat different phenotypic strengths (Fig 9B, D, F). The results of double knockdown were striking. Gap number increased significantly relative to either single knockdown (Fig 9C, E, G; quantified in H-J)—this was true both with the two weaker *RN-tre* shRNAs and the stronger one. Further, while RN-tre knockdown alone only led to gaps in the proximal testis, in double knockdowns gap number and extent were also increased distally (Fig 9B, D, F, cyna arrows). To verify these differences in effect along the proximal-distal axis, we computationally unrolled the testes, and manually highlighted gaps (Fig 9K-M). Ncad knockdown led to small gaps all along the proximal-distal axis (Fig 9K). In contrast, gaps seen after RN-tre knockdown were confined to the proximal region (Fig 9L). After double knockdown, large gaps were present both proximally and distally (Fig 9M). This was also true in the double knockdown using the weaker RN-tre line, where the effect on testis shape was less drastic (Fig 9G, cyan arrows), These data are consistent with RN-tre playing a role in adhesion parallel to that of Ncad. They suggest Ncad is sufficient for adhesion distally, even when RN-tre is knocked down, but if Ncad is knocked down RN-tre becomes important for maintaining adhesion distally as well. We also explored the effects of double knockdown of Ncad and Par-1, but in this case double knockdowns retained the very strong phenotype of Par-1 (Fig 9 N vs O), and thus we did not perform gap quantification.

**Fig 9.**
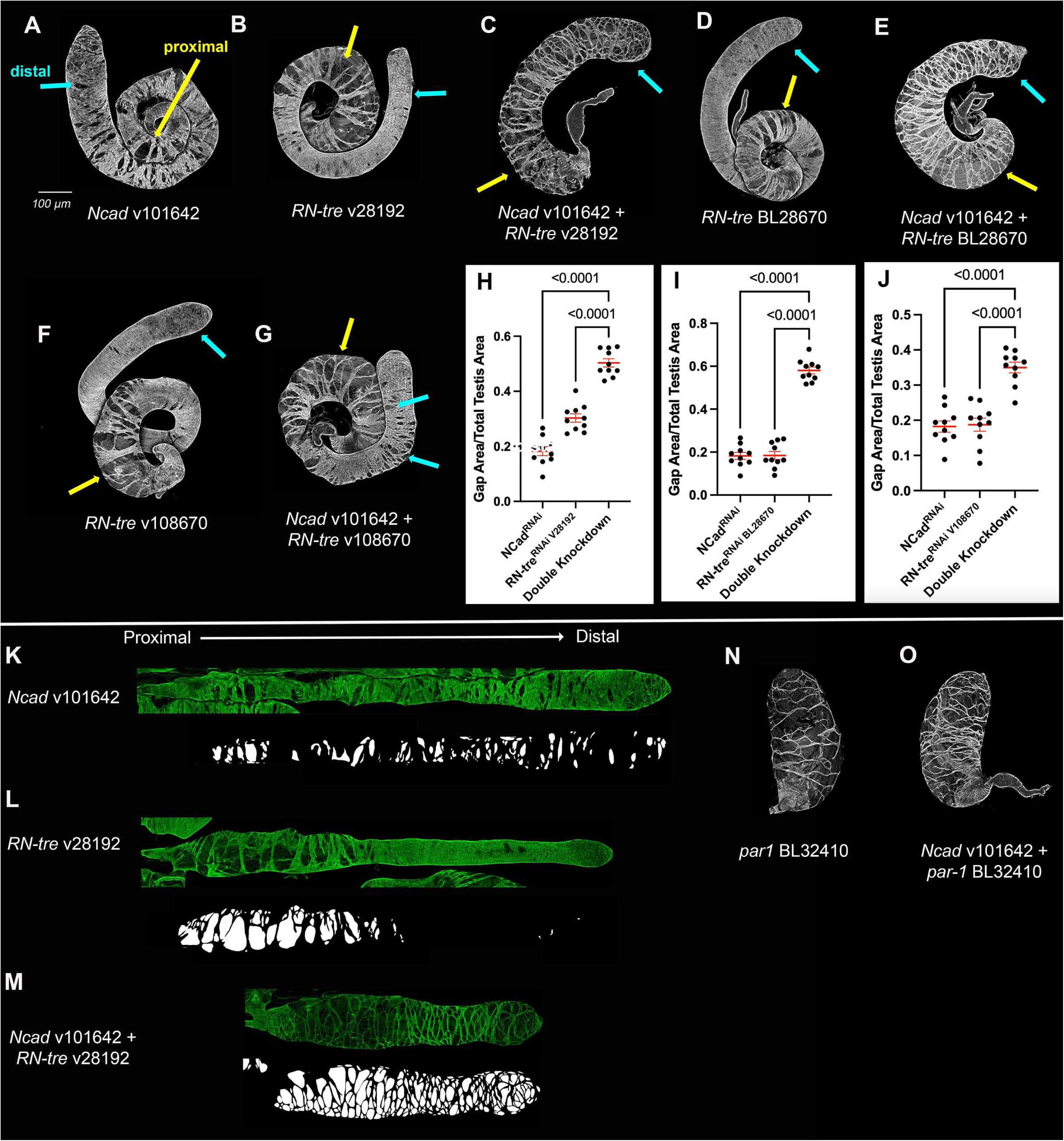
Double knockdown suggests broader roles for RN-tre in adhesion. **(A-G, L-M)** Testis from wildtype or adults expressing indicated shRNAs or mutant constructs. All are stained with phalloidin to reveal F-actin, highlighting muscles. shRNA lines or genotypes are indicated. All single knockdowns carry an additional UAS-driven gene to account for the fact that GAL4 may be limiting. Proximal=yellow arrows. Distal = cyan arrows. (A) Ncad knockdown alone. Moderate gaps all along the proximal-distal axis. (B,D,F) RN-tre knockdown with three different shRNAs. All result in moderate gaps restricted proximally (yellow vs cyan arrows). (C, E) Two of the Ncad RN-tre double knockdowns drastically elevate area covered by gaps, both proximally and distally. (G) The third double knockdown enhances gap area but is less drastic. **(H-J)** Quantification of Gap Area/Total Testis area, confirming the enhancement after double knockdown. The line represents the mean and error bars represent SEM, and all datapoints are indicated. **(K-M)** Computationally unrolled testis. K. Ncad knockdown. Small gaps all along the proximal-distal axis. L. RN-tre knockdown. Gaps are restricted to the proximal region. M. Ncad RN-Tre double knockdown. Large gaps are seen all along the the proximal-distal axis. (N) Par1 knockdown. (O) Ncad Par-1 double knockdown resembles Par-1 knockdown alone.

### Ecad and Ncad work in parallel to maintain adhesion

We entered these experiments thinking Ncad was the sole classical cadherin at work during myotube collective migration and testis morphogenesis. Consistent with this, mRNA levels of Ecad appeared lower than those of Ncad during and after migration, and we could not detect Ecad at cell junctions of migrating myotubes (Bischoff *et al*., 2025a; Bischoff *et al*., 2025b). However, during the screen we found that two different shRNAs targeting Ecad that led to mild to moderate defects in distal muscle coverage and shaping. We were intrigued by this role and wondered if the two classical cadherins might partially compensate for one another. To test this, we used shRNAs to knockdown both Ncad and Ecad.

Ncad knockdown led to frequent small gaps all along the proximal-distal axis (Fig 10 A vs B, arrows). The weaker of the two Ecad knockdown lines (BL32094) had only subtle distal defects (Fig 10C, arrow). After double knockdown with this *shg* shRNA, double knockdown testis largely resembled single Ncad knockdown, with small gaps all along the proximal-distal axis (Fig 10D, yellow arrows), with no increase in gap area (Fig 10I), and distal defects similar to or slightly stronger than those after Ecad knockdown (Fig 10D, cyan arrow). On its own, the stronger of the two *Ecad (shg)* shRNAs (v27081) also only had small gaps in distal muscle coverage (Fig 10E, cyan arrow). However, in the double knockdown with this shRNA, gap area was significantly increased relative to both single knockdowns (Fig 10J). In addition, double knockdown led to substantial defects in testis shaping, with testis diameter becoming extremely variable (Fig 10B vs F-H; n=10 testis/genotype). Thus, despite accumulating lower levels, Ecad does contribute to maintaining cell-cell adhesion during morphogenesis. The single knockdown phenotypes suggest Ecad is more important distally, but in Ncad’s absence Ecad plays roles all along the proximal-distal axis.

**Fig 10.**
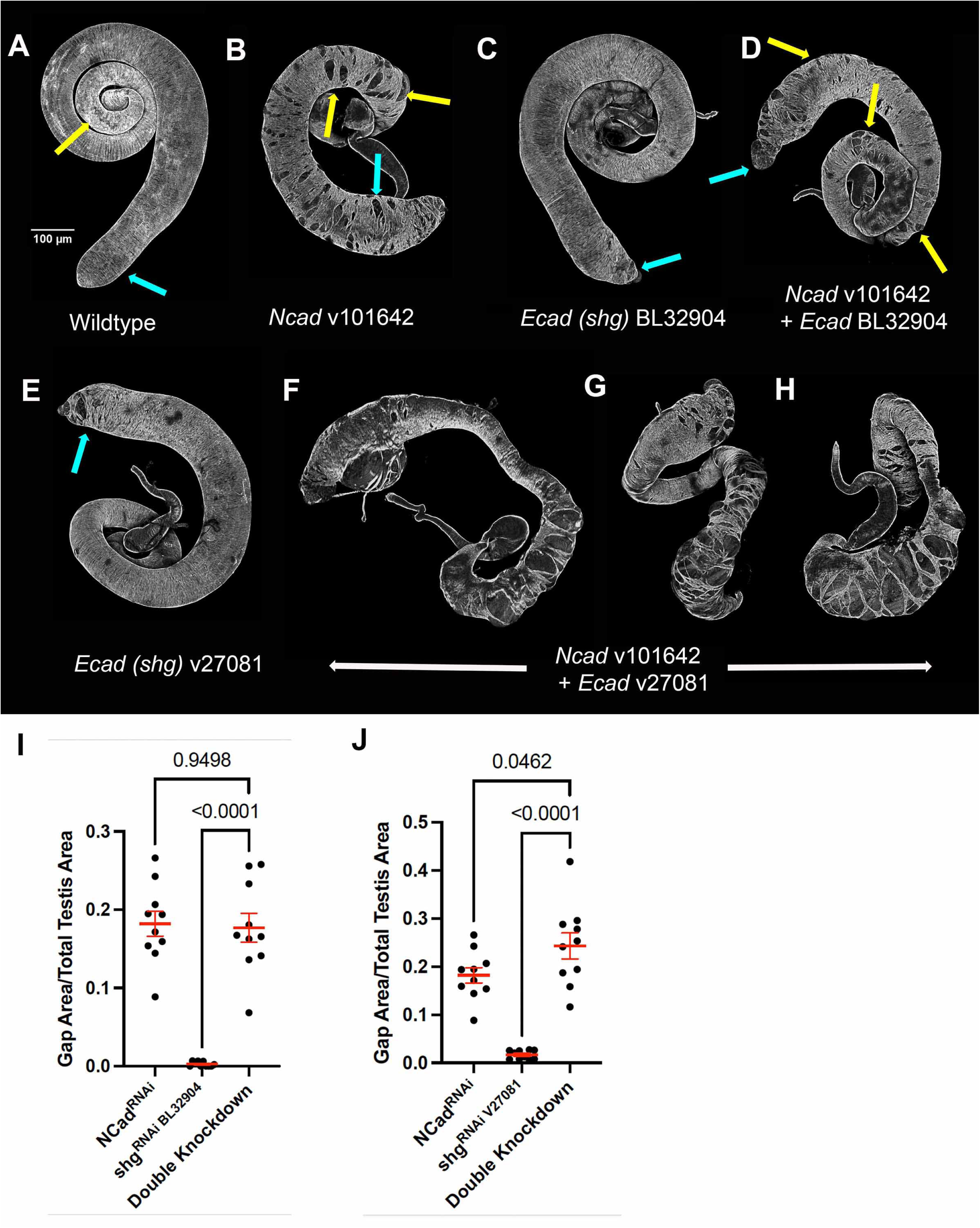
Ncad and Ecad work in parallel to maintain testis adhesion. **(A-H)** Testis from wildtype adults and mutant adults. All are stained with phalloidin to reveal F-actin, highlighting muscles. Proximal = yellow arrows. Distal = cyan arrows. (A) Wildtype. (B) Ncad knockdown. (C, E) Ecad knockdowns. V27081 has stronger, more penetrant distal defects. (D) The Ncad Ecad double knockdown with the weaker Ecad shRNA looks similar to the single Ncad knockdown. (F-H) The Ncad Ecad double knockdown with the stronger Ecad shRNA has elevated gap area and strong effects on testis shaping. **(I-J)** Quantification of Gap Area/Total Testis Area. Line represents the mean and error bars represent SEM. All datapoints are shown.

## Discussion

While animal bodies contain a diverse array of tissue types with distinct architectures and cell behaviors, cell adhesion mediated by classical cadherins is foundational for most. These differences are often simplified by characterizing tissues as epithelial versus mesenchymal, with the former more static and the latter often migratory. However, this is a continuum with different cell types positioned across the epithelial-mesenchymal spectrum(Bischoff and Mayor, 2025; Zhang *et al*., 2025). One key question for our field is to define the underlying molecular mechanisms that drive differences between tissues. Differential expression of different classical cadherins can be one driving force, with an Ecad/Ncad switch often seen as key to the epithelial-mesenchymal transition (EMT; (Loh *et al*., 2019)). However, some tissues express both cadherins, suggesting other contributors. One possibility is that different tissues make use of different subsets of the large array of proteins associating at the cytoplasmic face of cell-cell junctions. Here we use a model mesenchymal migrating tissue, the Drosophila testis nascent myotubes, to begin to define the junctional proteins important for myotube migration and testis morphogenesis.

### This mesenchymal tissue uses two different Classical cadherins

The simple model of EMT suggests that mesenchymal tissues derive their behaviors from the expression of Ncad versus Ecad. When we began our work, this mechanism appeared to be in action during testis morphogenesis. Ncad accumulates at cell junctions of myotubes and is important for their function. In contrast, immunofluorescence analysis suggested that Ecad accumulated at lower levels, so that we could not detect it at myotube cell junctions, while it was detectable in the overlying pigment cells (Bischoff *et al*., 2025a; Bischoff *et al*., 2025b). However, our current shRNA knockdown data suggest this assumption was an over-simplification. Two different shRNAs targeting Ecad led to mild to moderate defects in distal muscle coverage and shaping, suggesting that Ecad might play a modest role in migration or maintaining adhesion distally, unlike the broader role of Ncad in adhesion all along the proximal-distal axis. This hypothesis was further strengthened by double knockdown of Ncad and Ecad. This led to additive or synergistic effects, with elevated muscle gaps suggesting reduced adhesion all along the proximal-distal axis, and more dramatic effects on testis shaping. This could reflect temporally distinct roles for the two classical cadherins, with, for example, Ecad more important at later steps in the complex shaping of this organ. Alternately, Ecad may simply serve as a backup, reinforcing Ncad adhesion at times or places where stresses on cell junctions are maximal. Future work will be needed to discriminate between these models. It is worth noting that the Drosophila ovarian somatic border cells depend on Ecad rather than Ncad (Niewiadomska *et al*., 1999), and migrate in a more epithelial fashion.

### Migrating myotubes use a distinct subset of junctional proteins

Cadherin-catenin complexes organize complex arrays of dozens of proteins by multivalent interactions (Rouaud *et al*., 2020). Another way different tissues could exhibit differential architectures and behaviors is by using different subsets of these junctional proteins. Substantial information is available about the function of many of these proteins in epithelial cells in flies and mammals, but less is known about which junctional proteins act in mesenchymal cells. To begin to address this problem, we used three resources: a proximity proteomics screen identifying AJ proteins in the mammalian epithelial model cell line, MDCK cells (Choi *et al*., 2025), an RNAseq database of genes expressed in migrating Drosophila myotubes (Bischoff *et al*., 2025b), and genome-wide libraries of shRNAs targeting Drosophila genes (Dietzl *et al*., 2007; Ni *et al*., 2011). This gave us a candidate list of potential AJ proteins we could test for function during testis morphogenesis.

As expected, the core catenins play roles, though, as we observed in embryonic epithelia, the role of p120 is more modest than it is in mammals. In contrast, several proteins that play key roles in embryonic epithelia play modest or no role in migrating myotubes. Using validated shRNAs and in some cases mutants, we found that Cno, Pyd, Baz and aPKC are likely to play at most modest roles during testis morphogenesis. Consistent with this, both Cno and Pyd are not detectable at cell junctions of migrating myotubes, while they accumulate at easily detectable levels in the adjacent epithelial seminal vesicle. Intriguingly, both Baz and the apical polarity complex play important roles in Drosophila border cell migration (Pinheiro and Montell, 2004), where Ecad is the relevant classical cadherin (Niewiadomska *et al*., 1999). RNAi screens in that cell type support the lack of a role for Cno, but data for Pyd are more equivocal (Aranjuez *et al*., 2012).

Another protein complex that plays important roles in a subset of Drosophila epithelia is the Magi/ASPP/RASSF8 complex. They play important roles in the complex cell shape changes that create the stereotyped organization of the Drosophila ommateum during pupal eye development (Langton *et al*., 2009) (Zaessinger *et al*., 2015). This is a tissue where both Ecad and Ncad play roles (Mirkovic and Mlodzik, 2006), as do Cno (Matsuo *et al*., 1997), Pyd (Seppa *et al*., 2008), and aPKC (Wang *et al*., 2018). Our analyses, in this case using null mutants, reveal that the Magi/ASPP/RASSF8 complex is also largely or completely dispensable for testis morphogenesis. This is even more intriguing as all three proteins play roles in border cell migration (Aranjuez *et al*., 2012; Hohne *et al*., 2025), consistent with a possible restriction of their roles to tissues where Ecad predominates. We also found no role in testis morphogenesis for the CD2AP homolog Cindr, which also plays important roles in the eye development (Johnson *et al*., 2008). Thus, many proteins that are important in epithelial tissues are dispensable or have only minor roles in this mesenchymal tissue.

### Screening candidate AJ proteins provides multiple new leads for exploring collective cell migration

Our proximity proteomics list provided us with 58 candidate genes whose potential roles in myotube migration and testis morphogenesis we explored (Table 1). From this list, we identified ten genes with strong defects in testis morphogenesis, and twelve genes with moderate to mild defects (Fig 5, 6). These defects altered testis morphogenesis in multiple ways. Two, Par-1 and Dlg5 knockdown, led to dramatic alterations in testis shape and loss of muscle coverage. α-catenin knockdown phenocopied Ncad, as expected, with gaps all along the proximal distal axis. In contrast, RN-tre knockdown led to a unique phenotype in which gaps in muscle coverage were only present proximally. Most of the other genes with moderate to strong defects only altered the distal testis, ranging in severity from complete loss of distal muscle coverage to mild defects in the shape of the distal testis. Dsh knockdown also had a unique phenotype, with characteristic defects in the testis spiral and muscle alignment defects. These were strikingly similar to the defects we had previously seen when knocking down the Wnt ligand Wnt4 and its receptor Fz2 (Bischoff *et al*., 2025b), consistent with Dsh’s role in the canonical and planar polarity Wnt pathways. Twenty genes had very mild, lower penetrance defects, with almost all affecting distal testis shaping (Suppl Fig 3). While these were outside the normal wildtype range, it remains to be seen if these are genes with genuine but mild roles, or these were shRNAs that are not full effective at knocking down genes, or the mild phenotype was due to off target effects. Knockdown of the remaining genes had no effect on testis morphogenesis. While we tried to choose shRNAs with verified effectiveness in somatic tissues, we cannot rule out role for these genes based our negative data, except in the cases where a molecular null allele was available.

Several of our hits also are important for other collective cell migration events. One of the best studied migration events in Drosophila is border cell migration, where somatic follicle cells migrate as a group between the nurse cells to the anterior end of the oocyte (Montell *et al*., 2012). Several hits in our screen are implicated in border cell migration, including Par-1 (McDonald *et al*., 2008), Dlg5 (Aranjuez *et al*., 2012), Scrib (Campanale *et al*., 2022), Ena (Gates *et al*., 2009), and RN-tre (Laflamme *et al*., 2012). However, other genes with known roles in border cell migration, like Baz (Pinheiro and Montell, 2004)and RASSF8 (Hohne *et al*., 2025), were negative in our screen. Several genes we tested have known roles in Drosophila hemocyte migration: some, like Fit1 and Ena, also had effects in the testis whereas others like Rhea/Talin did not (Moreira *et al*., 2013) (Tucker *et al*., 2011). We suspect that the different modes of collective cell migration may each rely on its own unique subset of junctional proteins.

### Par-1 and RN-tre play distinct roles in testis morphogenesis

We hope our screen will provide a starting point for the community to explore the mechanisms by which genes that scored as hits in our screen influence collective cell migration. As an illustration of how one might proceed, we examined two of our strongest hits, Par-1 and RN-tre. Both gave similar phenotypes with two or more shRNAs, reducing concerns about off-target effects. They had dramatically different adult phenotypes. Par-1 knockdown left the testis in its original oval shape, with small numbers of highly stretched muscles covering a small part of its surface. In contrast, RNA-tre knockdown did not disrupt testis elongation and was compatible with spiralization. However, gaps in muscle coverage were common, but these were restricted to the proximal testis, in contrast to most of our hits where distal defects were common.

As a first avenue to explore the mechanism by which these proteins affect testis morphogenesis, we turned to live-imaging the testis during and just after migration. To our surprise, RN-tre knockdown testis were indistinguishable from wildtype at these stages, with no delay in migration, and no alteration in gaps or gap closure. This suggests the morphogenetic process RN-tre affects occurs after the completion of migration, when spiralization changes the shape of the testis. In contrast, Par-1 knockdown significantly delayed migration. Par-1 knockdown testis had significantly larger gaps, suggesting defects in gap closure. We were intrigued by the fact that the adult phenotype of Par-1 knockdown was strikingly similar to that caused by knocking down cytoplasmic myosin heavy chain or regulatory light chain (Bischoff *et al*., 2021; Bischoff *et al*., 2025a). One role of myosin during myotube migration is to assist in gap closure, with actomyosin cables forming around gaps, similar to those seen in embryonic wound closure. When cytoplasmic myosin is knocked down, actomyosin cables are lost and gaps become larger and more frequent (Bischoff *et al*., 2025a). In these conditions, cells surrounding gaps extend more filopodia into them, as they do at the normal leading edge. Par-1 knockdown also increased filopodia on cell borders next to gaps, consistent with a reduction in actomyosin cable formation. Together, these data suggest Par-1 and myosin might work together in myotube migration. Relevant to this, Par-1 has a known role in border cell migration (McDonald *et al*., 2008). In that cell type Par-1 promotes myosin inactivation by phosphorylating and inactivating its negative regulator myosin phosphatase. It will be interesting to further explore whether Par-1 is acting by a similar mechanism in myotubes.

The phenotype of RN-tre knockdown was intriguing, as it led to small gaps suggestive of adhesion defects. However, unlike Ncad knockdown, which leads to adhesion defects all along the proximal-distal axis, RN-tre knockdown adhesion defects are restricted to the proximal testis. Further, Ncad knockdown led to noticeable defects during migration, including separation of individual cells from the collective and increased gap number (Bischoff *et al*., 2021), but our live imaging revealed no defects in migration after RN-tre knockdown. Instead, this suggested RN-Tre defects arise during spiralization. To begin to define the relationships between RN-Tre and Ncad in maintaining adhesion, we used double knockdown. The results were intriguing. Double knockdown animals had more and larger gaps. Most surprising, RN-tre/Ncad double knockdown increased gap number and size distally, where RN-Tre knockdown alone had no effect. We suspect RN-tre supports adhesion all along the proximal-distal axis, but its role distally is redundant when Ncad is fully functional. Our current efforts to understand the cellular events underlying spiralization will provide the foundation for defining the role RN-tre plays in later testis morphogenesis.

Exploration of RN-tre’s roles in other contexts provides diverse possibilities for molecular mechanisms. It’s role as a GAP for Rab5, Rab41, and Rab43 suggest roles in protein trafficking (Lanzetti *et al*., 2000; Haas *et al*., 2005). RN-tre has known roles in regulating integrin-based focal adhesion turnover and chemotactic migration in cultured cells (Palamidessi *et al*., 2013). It seems likely both cadherin and integrin trafficking play roles in testis morphogenesis. RN-tre also regulates trafficking and signaling of receptor tyrosine kinases (Lanzetti *et al*., 2000), and myotube migration requires the fibroblast growth factor receptor Heartless (Rothenbusch-Fender *et al*., 2017). Finally, evidence in cultured Drosophila cells suggests roles in regulating Rho and myosin contractility (Platenkamp *et al*., 2020). RN-tre was one of the genes that scored positive in an shRNA screen for RabGAPs with roles in border cell migration (Laflamme *et al*., 2012), though the authors did not pursue this hit further to explore mechanisms. Future work will help sort our which of these many roles is key in testis morphogenesis.

## Materials and Methods

### Fly Genetics

We performed crosses in a 25°C. We developed prepupae for live cell imaging at 26.5°C. All shRNA, gRNA, and null allele lines used can be found in Supplemental Table 2. For control crosses, we used w^1118^ flies and refer to them as “wildtype.” For the screen in the adult testis, we used the driver lines *mef2-Gal4* (BL27390) and *UAS-Cas9*; *mef2-Gal4*/TM6B (BL67075). For pupal testis staining and live cell imaging, we used the driver line *UAS-lifeact-GFP/*cyo; *mef2-Gal4,* UAS mCherry nls (Bischoff *et al*., 2021). For our double knockdown experiments in the adult testis, we used the driver line *N-cad shRNA^V101642^*/CyO; *mef2-Gal4/*MKRS. We also used *UAS-GFP; mef2-Gal4* to coexpress UAS-green fluorescent protein (GFP) in single-shRNA controls to include the same number of UAS-promoters in all conditions, correcting for potential competition for Gal4 binding.

### Adult Testis Staining and Microscopy

For the screen in the adult testes, samples were dissected 3-5 days post eclosion in 1.5x phosphate-buffered saline (PBS). The testes were then fixed for 20 minutes in 3.7% paraformaldehyde (PFA) in 1.5x PBS. Post fixation, testes were washed three times in 1.5x PBS with 0.1% Tween 20 (1.5x PBST). Testes were stained overnight in 1:500 Phalloidin (*Thermo Fisher, Alexa Flor 488)* in 1.5x PBST at 4°C. We then washed the samples three times in 1.5x PBS and mounted them on 35mm glass bottom dishes in PBS. To image them, we used an LSM Pascal (Zeiss) with a 10x (Zeiss EC Plan-Neofluar 10x/0.3) dry objective. We created maximum intensity projections of large z-stacks taken of the adult testes, and modified brightness and contrast using Photoshop (Adobe) and Fiji (ImageJ), making all structures visible.

We processed images of adult testis using the Fiji macro “Surface Peeler” (https://forum.image.sc/t/surface-peeler-fiji-macro-for-fast-near-real-time-3d-surface-identification-and-extraction/61966). This macro recognizes the upper surface and automatically deletes every pixel under a layer with defined thickness. (we used the following values: upper boundary 0 µm, lower boundary 4 µm). As a filter type we used Gaussian Blur, with a radius of 2 and Huang for thresholding. We modified the macro slightly, to reverse the stack order and to add an empty stack on top. Finally, we created a maximum intensity projection of the “surface-peeled” data and changed the contrast in Fiji to make all structures visible. This eliminates interference by brightly stained sperm inside the testis.

### Pupal Testis Staining and Microscopy

For the fixed pupal tissue images, animals were collected as prepupae, aged, and then testes were dissected at 32 hours APF or 40 hours APF. The testes were then fixed for 20 minutes in 3.7% PFA in 1.5x PBS. Post fixation, testes were washed three times in 1.5x PBS. Testes were then blocked in 1.5x PBST with 10% Normal Goat Serum for thirty minutes (ThermoFisher Scientific). Primary antibodies were then applied overnight at 4°C. We used the following antibodies: anti–N-cadherin (1:500; DSHB, DN-Ex #8), anti-arm (1:100; DSHB), anti-cno (1:1000; (Sawyer *et al*., 2009)), anti-pyd (1:100; DSHB), and anti-GFP (1:1000; Rockland) in 1.5x PBST with 10% Normal Goat Serum. The next day, testes were washed three times in 1.5x PBS and blocked a second time. Secondary antibodies were then applied at room temperature for two hours. We used Dylight secondary antibodies at 1:1000 (ThermoFisher Scientific) in 1.5x PBST with 10% Normal Goat Serum. Finally, testes were mounted in Fluoromount-G (ThermoFisher Scientific) and imaged using a Zeiss LSM 980 with Airyscan 2 and a 63x/1.4 oil Plan Apochromat objective.

### Live Cell Imaging

For the live cell imaging, samples were collected as prepupae and aged to 31-32 or 40 hours APF (Bischoff and Bogdan, 2023) Aged pupae were dissected in M3 Medium (Shields and Sang, Sigma-Aldrich) with 10% FCS (ThermoFisher Scientific) and 1x penicillin/streptomycin. The live testes were mounted in a glass bottom dish and encased in 0.5% low gelling agarose (Sigma-Aldrich) in M3. We imaged samples using a Nikon Ti2 inverted microscope with a Yokogawa CSU-W1 spinning disc and Hamamatsu ORCA-fusionBT sCMOS camera and a 25x/1.05 Silicone Apochromat objective. We modified brightness and contrast using Photoshop (Adobe) and Fiji (ImageJ), making all structures visible.

### Analysis and Quantifications

For quantification distance migrated at 32 hours APF in the live cell imaging stills, we measured distance from the bottom of the migrating sheet to the front of the migrating sheet using the “line” tool in Fiji (ImageJ). We used a machine learning-based classification approach to quantify area of filopodia/cell edge length (as a proxy for filopodia density), gap-area and cell area in 32 h APF pupae. To this end, we used the “Trainable Weka Segmentation” within Fiji (Arganda-Carreras *et al*., 2017). We created a training dataset that did not include any of the data we used for the quantification and trained Weka to recognize the cell body (excluding filopodia), filopodia and cell gaps as individual features. In preparation for quantification, we created rectangular region of interests (ROIs) of maximum-projected and denoised still images of 32 h APF pupae with Lifact-GFP (green channel) and cherry-nls (red channel) expressed in TNMs, using (Nikon Denoise.ai software). The trained model was then applied to the experimental data and the labeled output image (“Create Result”) was used for further analysis. We subsequently processed the resulting labeled images using a custom Fiji Macro that uses the Fiji Plugin MorphoLibJ (Legland *et al*., 2016). Cell gaps were smoothed by morphological closing (radius = 2) and binarized prior to particle analysis to obtain gap perimeter and area. The script subtracts image border-associated perimeter contributions to avoid edge artifacts. Further, the script excludes the upper-most gap from the gap area quantification as this represents the area in front of the cell sheet. It’s important to note that this can cause artifacts, when gaps within the sheet are interconnected and open toward the front, as was seen after *sqh*-RNAi. Filopodia and cell body classes were independently binarized and quantified by particle analysis to determine total filopodia and cell area per image. The ratio between total filopodia area (in µm²) and gap perimeter (free edge length in µm) was then calculated as a quantitative readout of filopodial density. The script creates a .csv-file as an output. Cell area per cell was calculated by manually counting the number of cells within each ROI and dividing the total cell area by this value.

For quantification of the adult testes double knockdowns, we manually traced gaps in the muscle sheet using Fiji (ImageJ). We then traced an outline of the whole testis using the “polygon” tool and calculated the percentage of gaps relative to the whole testis using Excel. For the analysis of proximodistal gap distribution after single and double knockdown, we followed the protocol described in (Bischoff *et al*., 2025a). Briefly, we manually traced gaps on the image and changed the color of the gap-ROIs to white. We used the “segmented line” tool to define the proximodistal axis and then linearized the testes using “Edit>Selection>Straighten” with a width of 300 μm. We linearized the imaging data and the “gaps-in-black-and-white” channel.

For statistical analysis of the live cell imaging stills and adult testes double knockdowns, we used Welch’s t-test or One Way ANOVA with multiple comparisons. We plotted data showing mean and SEM error bars in GraphPad Prism.

## Author Contributions

This project was conceived by Sarah Clark, Maik Bischoff and Mark Peifer. Sarah Clark led the team who carried out the screen. Siobhan Morris also played a substantial role in the screen, while Joseph Dordor1, Lawrencia Amo, and Taino Encarnacion helped in early stages. Sarah Clark did the live imaging, with analysis provided by Maik Bischoff. Sarah Clark and Siobhan Morris stained pupal testis., Renick Wiltshire validated shRNAs in the embryo. Sarah Clark, Maik Bischoff and Mark Peifer drafted the manuscript with editing and comments by the other authors.

## Acknowledgements

We would like to thank Flybase for the tremendous resource they provide for assessing previous use of different shRNA lines, the Bloomington and Vienna *Drosophila* stock centers for providing many, many fly stocks, the Dvelopmental Studies Hybridoma Bank for antibodies, Dr. Nathanaël Prunet and the UNC Biology Department Imaging Core for imaging advice, Dr. Jocelyn McDonald for providing antibodies and a valuable discussion, Corbin Jones for advice on RNAseq data, Kevin Slep and Dan Bergstralh for comments on the manuscript, and the Peifer, Slep, Bergstralh/Finegan, Williams, and Lovegrove labs for feedback throughout. Work in the Peifer lab is supported by NIH R35 GM118096.

---

**Suppl Fig 1.**
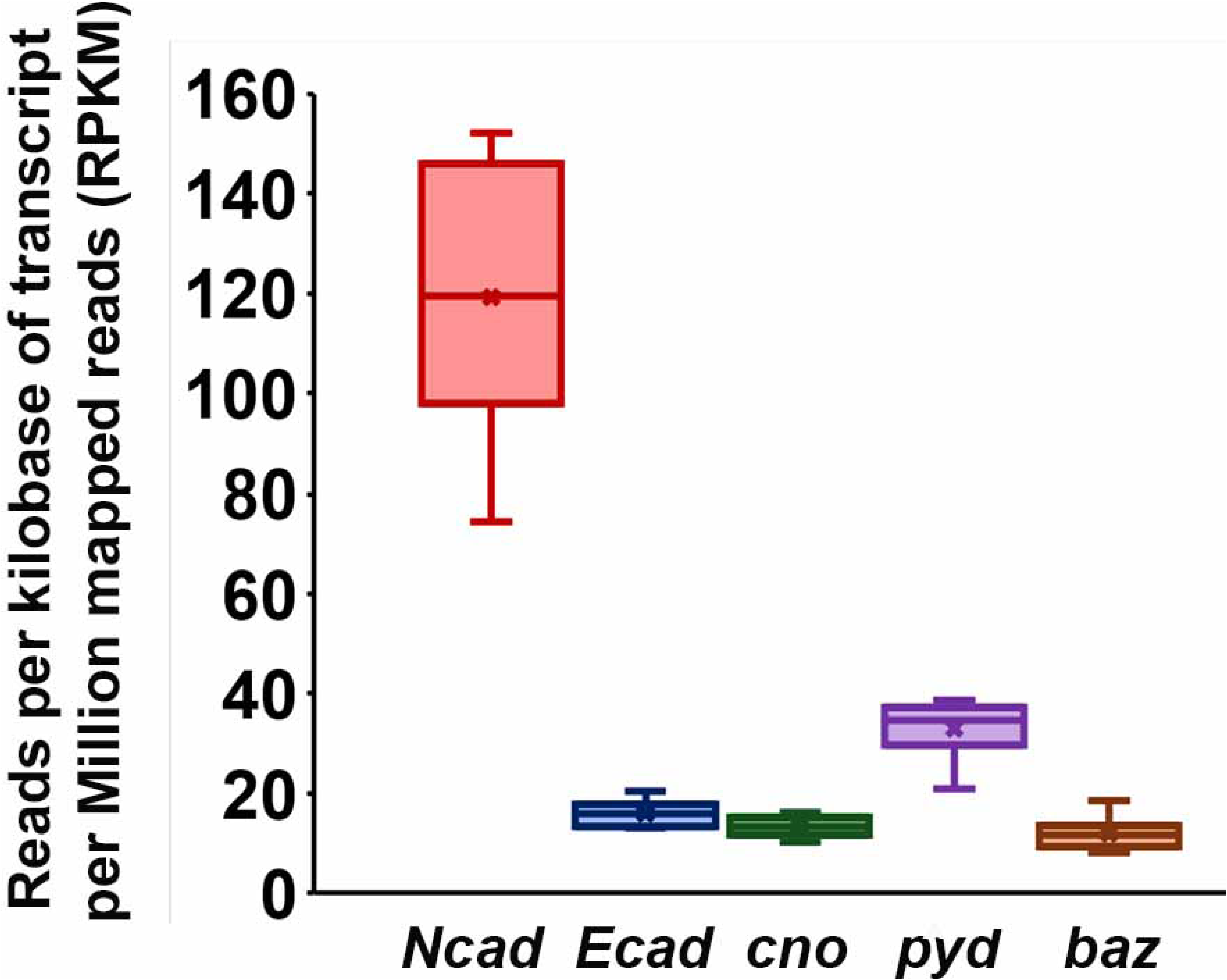
RNAseq data of classical cadherins and some key epithelial regulators. Box and Whisker plot of the RNAs detected during migration (31 hour timepoint) from the RNAseq experiment published in Bischoff et al. (2025). They represent the data from the six replicates of each gene. The box includes the first to the third quartiles, while the whiskers are the minimum and maximum values. The line indicates the median value and the X the mean value.

**Suppl Fig 2.**
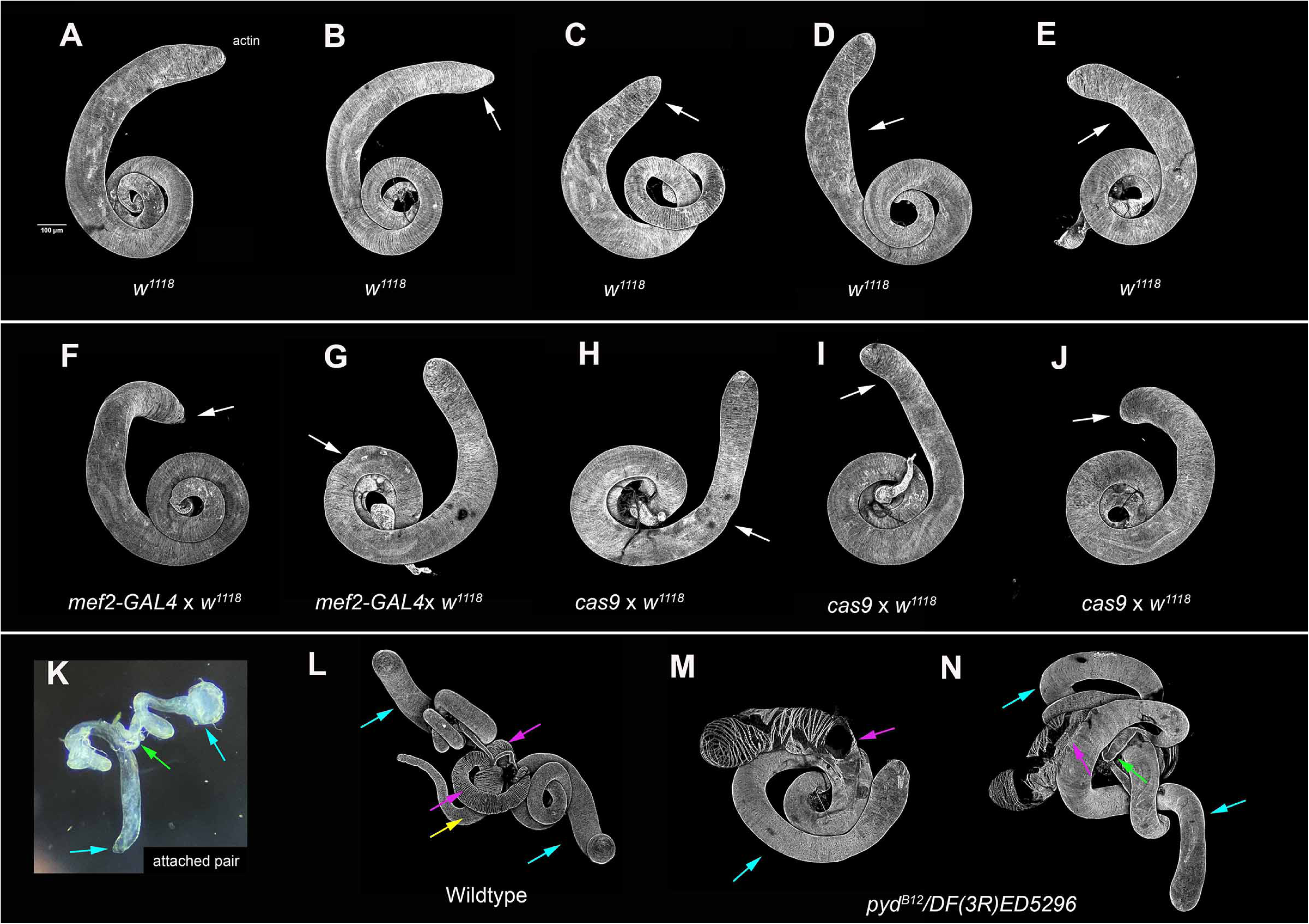
The phenotypic range of wildtype testes. All are stained with fluorescently-labeled phalloidin to reveal F-actin, highlighting muscles. **(A)** Typical wildtype testis. **(B-N)** Examples of the most severe deviations in testis shape seen in testes from wildtype adults or controls for use of the *mef2-*GAL4 driver or the *cas9* line. White arrows denote mild variation in adult testis morphogenesis. Gaps in muscle coverage were never observed. Even minor defects like these were seen in <10% of the testis from a wildtype or control genotype. (A-E) Wildtype *(w^1118^*) natural variation. (F-G) *mef2-Gal4* crossed with *w^1118^*natural variation. (H-J) *Cas9* crossed with *w^1118^* flies natural variation. (K) Low magnification view of a knockdown adult testis pair with divergent phenotypes. The testis pair is attached by the adult seminal vesicle (attachment shown at green arrow). The two testes have different distal testis phenotypes. The cyan arrow on the left points to a wildtype-like distal testis. The cyan arrow on the right points to an adult testis with a moderate distal muscle coverage defect. (L) Whole adult *Drosophila* reproductive system: wildtype adult testes, paragonia, and genital tract. Cyan arrows point to wildtype adult testes. Magenta arrows point to the two paragonia. The yellow arrow points to the genital tract. (M, N) *pyd^B12^/ DF(3R)ED5296* adults have a testis-paragonia fusion. Cyan arrows point to the adult testes. Magenta arrows point to the areas of the testes that are fused with the paragonia. In N, the green arrow points to the attachment of the adult testes to one another.

**Suppl Fig 3.**
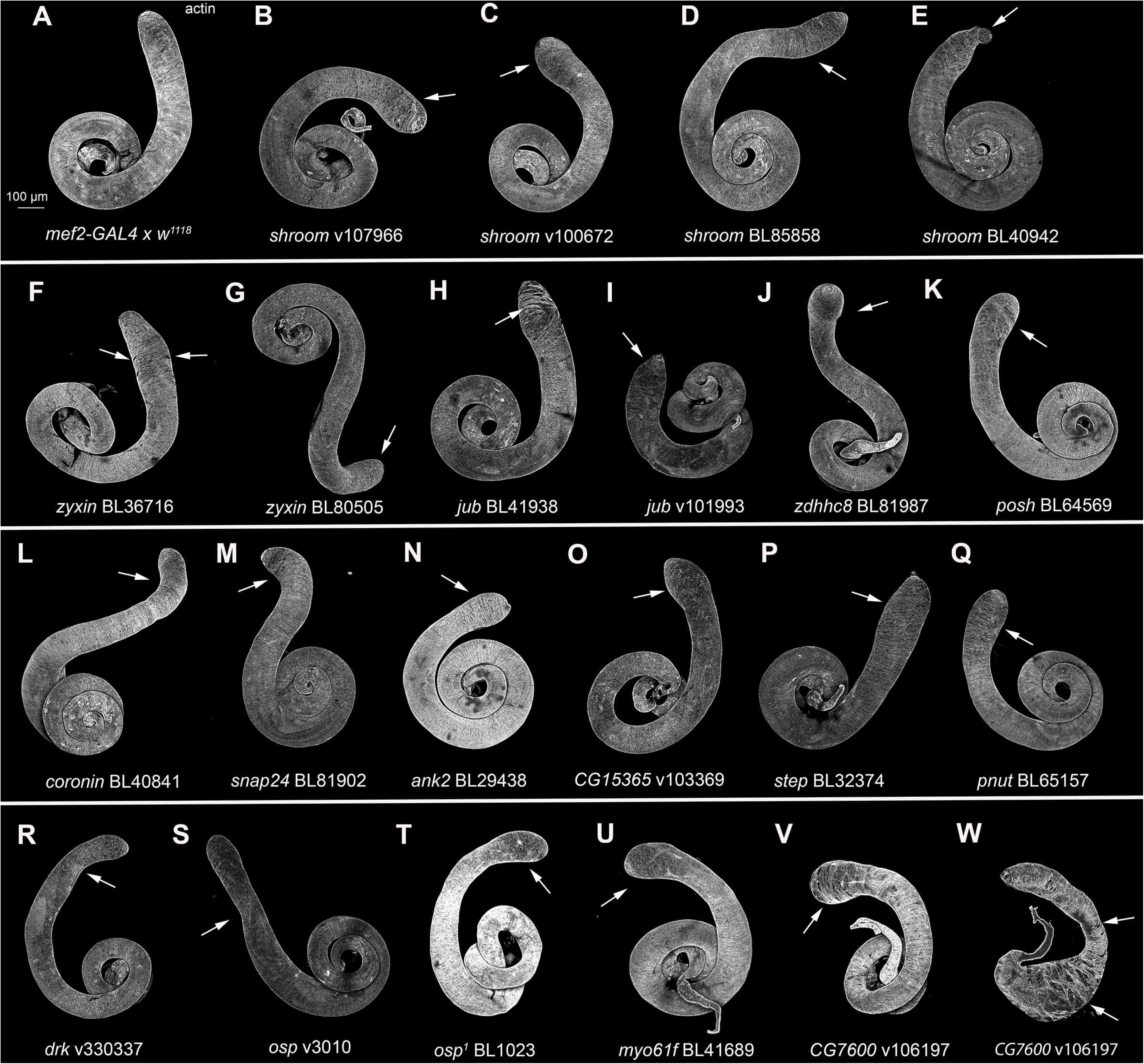
Some gene knockdowns mildly impact testis morphogenesis. Testis from wildtype, adults expressing indicated shRNAs, or homozygous mutants. All are stained with phalloidin to reveal F-actin, highlighting muscles. shRNA lines or genotypes are indicated. Arrows denote defects in adult testis morphogenesis. **(A)** Wildtype. **(B-E)** Shroom (Shrm) regulates contractility via effects on Rok and myosin. We observed phenotypes with four *shrm* shRNAs, with variable effects on distal testis shape. Two led to mild distal testis broadening with moderate penetrance (v107966 10/17 testes, v100672 9/23 testes). Two other shRNAs had lower penetrance phenotypes leading to a flipped distal tip or a pointed distal testis (BL85858 9/40 testes, BL40942 2/17 testes). (**F, G)** We tested two shRNAs targeting Zyxin, a LIM domain protein with roles in both cell-cell and cell-matrix junctions. One gRNA had mild, low penetrance defects in distal muscle coverage and shaping (BL36716 7/32 testes) while the other shRNA had a mild flipped distal testis (BL80505 5/18 testes). **(H, I)** The Ajuba LIM protein (Jub) localizes to AJs under tension. We tested four *jub* shRNAs, of which two had a phenotype. One gave a mild distal muscle coverage phenotype (BL41938 8/17 testes) while another gave a mild distal testis broadening phenotype (v101993 8/18 testes). **(J)** Zinc finger DHHC-type containing 8 (Zdhhc8) is a protein S-acyltransferase that has been implicated in regulating tissue growth through Scrib. We observed mild shaping defects including a flipped distal tip and low penetrance distal testis broadening in one of two gRNA/shRNAs tested (gRNA BL81987 12/21 testes). **(K)** Knockdown of Posh, a scaffolding protein that regulates TNF-JNK signaling, showed a mild broadening and low penetrance flipped distal tip phenotype (BL64569 7/20 testes). **(L)** Coronin regulates the stability of actin filaments. We tested two *coronin* shRNAs, one of which had a mild flipped distal testis phenotype (BL40841 6/18 testes), while the other caused embryonic lethality. **(M)** Synaptosomal-associated protein 24kDA (Snap24) regulates vesicle trafficking and exocytosis. We tested three *snap24* shRNA/gRNAs, one of which had a mild flipped distal testis phenotype (BL81902 5/20 testes). **(N)** Ankyrin 2 (Ank2) is a membrane-cytoskeleton linker with known roles at synapses. We observed a low penetrance mild broadening of the distal testis in the single shRNA we tested (BL29438 4/20 testes). **(O)** CG15365 encodes the single Drosophila member of the LZTS family; LZTS2 acts as an alpha-catenin effector (Wang *et al*., 2025). One of two shRNAs targeting CG15365 led to mild broadening of the distal testis with moderate penetrance (v103369 10/24 testes). **(P)** Mild enlargement of the distal testis was observed with an shRNA targeting the ArfGEF Steppke (Step; BL32374 9/41 testes). **(Q)** Mild enlargement of the distal testis was observed with an shRNA targeting the septin Pnut (BL65157 4/19 testes). **(R)** Knockdown of the receptor tyrosine kinase adapter Downstream of receptor kinase (Drk; the GRB2 ortholog) led to mild distal shaping defects (V330337 9/18 testes). **(S-T)** Outspread (osp), the homolog of MPRIP (Myosin Phosphatase Rho-Interacting Protein), acts as an adaptor to target myosin light chain phosphatase to the actin cytoskeleton. shRNA knockdown led to mild distal shaping defects (v3010 5/16 testes). We also tested a viable hypomorphic allele of *osp*, *osp^1^,* which we observed to have a mild broadening of the distal testis (4/18 testes). **(U)** Myosin 61F (Myo61F) is an unconventional myosin. When Myo61F was knocked down in the testis, it led to mild distal testis broadening with a low penetrance flipped distal tip (BL41689 6/20 testes). **(V-W)** CG7600 knockdown had a variable phenotype. B-E and J-W are included in our Class 4 (testis shaping defects). F-I are included in our Class 3 (distal muscle coverage defects).

